# Ciliopathy-associated protein, CEP290, is required for ciliary necklace and outer segment membrane formation in retinal photoreceptors

**DOI:** 10.1101/2025.01.20.633784

**Authors:** Abigail R. Moye, Michael A. Robichaux, Melina A. Agosto, Carlo Rivolta, Alexandre P. Moulin, Theodore G. Wensel

## Abstract

The most common genetic cause of the childhood blinding disease Leber Congenital Amaurosis is mutation of the ciliopathy gene *CEP290*. Though studied extensively, the photoreceptor-specific roles of CEP290 remain unclear. Using advanced microscopy techniques, we investigated the sub-ciliary localization of CEP290 and its role in mouse photoreceptors during development. CEP290 was found throughout the connecting cilium between the microtubules and membrane, with nine-fold symmetry. In the absence of CEP290 ciliogenesis occurs, but the connecting cilium membrane is aberrant, and sub-structures, such as the ciliary necklace and Y-links, are defective or absent throughout the mid to distal connecting cilium. Transition zone proteins AHI1 and NPHP1 were abnormally restricted to the proximal connecting cilium in the absence of CEP290, while others like NPHP8 and CEP89 were unaffected. Although outer segment disc formation is inhibited in CEP290 mutant retina, we observed large numbers of extracellular vesicles. These results suggest roles for CEP290 in ciliary membrane structure, outer segment disc formation and photoreceptor-specific spatial distribution of a subset of transition zone proteins, which collectively lead to failure of outer segment formation and photoreceptor degeneration.

## Introduction

Primary cilia are thin (∼300 nm diameter), signaling hubs that protrude from almost every eukaryotic cell at some point during the cell cycle (Mill et al., 2023, Ishikawa et al., 2021, Wensel et al., 2021). Like motile cilia, primary cilia have a central bundle of 9 doublet microtubules (DMT), termed the axoneme, which extends distally from the mother centriole of the centriole pair making up the basal body (BB). Along the length of the cilium, axonemal doublet microtubules (MTs) gradually transition to singlet microtubules, whose number declines as the MTs terminate (Sun et al., 2019, Kiesel et al., 2020, Moye, 2018, Ott et al., 2023). Surrounding the MTs is a ciliary membrane, which contains a specialized repertoire of membrane channels and receptors (Pazour et al., 2002, Pazour and Witman, 2003, Dutta and Ray, 2022, Garcia et al., 2018, Rohatgi and Snell, 2010). Defects in multiple genes encoding cilium-associated proteins lead to disruption of the normal spatial distribution of ciliary components and loss of ciliary structure and function (Horani and Ferkol, 2021, Van De Weghe et al., 2022, McConnachie et al., 2021).

Primary cilia are essential structural organelles in many sensory neurons, including photoreceptor neurons of the vertebrate retina, whose light-sensing compartment, the outer segment (OS), is a modified primary cilium (Wensel et al., 2021). In the OS, the conserved 9+0 DMTs serve as a structural support and trafficking scaffold for the formation of the OS membranous discs, which are packed with phototransduction proteins, including rhodopsin, the G-protein, transducin, and cGMP-specific phosphodiesterase-6 (Wensel et al., 2021, Liu et al., 2003). The OS cilium is a distinct compartment, separated from the biosynthetic photoreceptor inner segment by the bridge-like connecting cilium (CC). The CC, approximately 1100 nm long and 300 nm in diameter (Potter et al., 2021a, Gilliam et al., 2012), is thought to be analogous to the transition zone (TZ), a 200 to 300 nm long region present in all cilia between the distal BB and proximal axoneme (reviewed in (Wensel et al., 2021, Mercey et al., 2024)). As with other TZ, the CC connects the BB to the axoneme, and, with the exception of some CC-specific proteins such as the retina-specific isoform of RPGR (Hong et al., 2003, Patnaik et al., 2015), there is considerable overlap of protein components and structural features between CC and other TZ. The conserved structural features include the Y-links, filamentous structures which yoke the DMTs to the ciliary membrane; and the ciliary necklace, a “beads on a string” transmembrane structure that sits along the external face of the ciliary membrane (Insinna et al., 2008, Brown et al., 1963, Knabe and Kuhn, 1997, Steinberg and Wood, 1975, Wensel et al., 2016, Sun et al., 2019, Potter et al., 2021b, Ringo, 1967, Gilula and Satir, 1972, Robichaux et al., 2019, Zhang et al., 2024). The molecular composition of these structures is unknown, but they have been proposed to regulate trafficking into and out of the cilium and to aid in providing structural integrity (Muresan and Besharse, 1994, Garcia-Gonzalo and Reiter, 2017, Pedersen et al., 2012).

Although the photoreceptor CC contains many of the proteins and structural components observed in the TZ of other primary cilia and flagella, including ciliary transport proteins like the intraflagellar transport (IFT) proteins and kinesin motors (Robichaux et al., 2019, Insinna et al., 2008), there are a number of features, such as spatial distributions of certain cilium-associated proteins, that differ between photoreceptor CC and TZ of cilia in most other cell types. For example, Centrosomal Protein 290kDa (CEP290) is confined to a region at the base of the TZ in many primary cilia, but is found throughout the CC in rod cells (Potter et al., 2021b), and centrins, small Ca^2+^ binding proteins, generally localize to the BB in primary cilia, but occupy the lumen of the CC axoneme throughout its length in rods (Uytingco et al., 2019, Robichaux et al., 2019, Chen et al., 2024). In contrast, other TZ proteins such as CEP78 and NPHP8/RPGRIP1L only localize at the proximal end of the CC (PCC) in photoreceptors (Potter et al., 2021b, Nikopoulos et al., 2016). Ablation of another LCA-associated ciliary gene, spermatogenesis associated 7 (*Spata7*), results in redistribution of several proteins from the mid- and distal-CC to its base but does not affect the distribution of other TZ proteins (Dharmat et al., 2018). These results suggest there may be distinct ciliary compartments within the CC itself.

Mutations in ciliary genes often result in multi-syndromic diseases termed ciliopathies (Mitchison and Valente, 2017, Reiter and Leroux, 2017, Van De Weghe et al., 2022), which have pleiotropic phenotypes that affect the brain, organ laterality, kidney, lungs/trachea, skeleton, muscles, ear, or eyes. Retinal degeneration leading to blindness is a common symptom of a number of multi-syndromic ciliopathies, but it can also occur as an isolated “non-syndromic” disease as a consequence of certain mutations in ciliopathy genes (Goyal and Vanita, 2022, Murphy et al., 2015, Riazuddin et al., 2010, Fujita and Swaroop, 1996, Meindl et al., 1996, Littink et al., 2010, den Hollander et al., 2008). This phenotypic variability highlights the exceptional importance of cilia in photoreceptor function and retinal health. Mutations in CEP290, a core TZ protein, are the leading cause of the severe blinding disease, Leber Congenital Amaurosis (LCA), in which patients lose vision as early as 2 years of age (Tsang and Sharma, 2018, den Hollander et al., 2006). In addition, some *CEP290* mutations can cause non-syndromic retinitis pigmentosa (Birtel et al., 2018) or multi-syndromic disorders with associated retinal degeneration, such as Bardet-Biedl Syndrome and Joubert Syndrome (Radha Rama Devi et al., 2020, Sayer et al., 2006, Valente et al., 2006, Coppieters et al., 2010, Baala et al., 2007, Leitch et al., 2008, Frank et al., 2008, Brancati et al., 2007). In primary and motile cilia models, loss of CEP290 has been shown to cause complete disruption of ciliogenesis (Conkar et al., 2017, Kim et al., 2008, Tsang et al., 2008, Shimada et al., 2017) or defects in ciliary extension and trafficking. The mechanisms behind these defects remain poorly understood, as do the photoreceptor specific functions of CEP290.

Previously, using super-resolution microscopy and electron microscopy, we localized CEP290 along the length of the CC in photoreceptor cells and characterized the CC morphology in three different CEP290 mutant mouse models, determining that CEP290 mutations caused a decrease in CC diameter, but overall structure appeared relatively normal (Potter et al., 2021b). To further explore the role of CEP290 and CEP290 defects in photoreceptor CC/OS development and compartmentalization, and to assess more thoroughly the structural aberrations of the photoreceptor sensory cilium caused by CEP290 mutations, we have now performed an in-depth examination of the structure of the photoreceptors at various timepoints throughout photoreceptor ciliogenesis, including post-natal day 10 (P10), when CC are fully formed, OS have started forming, and photoreceptor cell death in the CEP290 mutants is not extensive, as well as earlier time points (P3 and P7) when cilia are just emerging.

For this study, we used a complete knockout (CEP290^KO^) and a C-terminally truncated mutant, which we have termed near-null, or CEP290^NN^ (Cep290^tm1.1Jgg^/J) in addition to WT mice. Both mouse models are whole-body mutants, phenotypic of Joubert Syndrome. In the CEP290^NN^ mouse model, exons 37 and 38 within the myosin-tail homology domain (domain of CEP290 reported to interact with the ciliary membrane (Drivas et al., 2013)) are removed, creating an early STOP codon and prompting nonsense-mediated decay. However, low levels of truncated CEP290 protein (∼200kDa) are still generated, with disruption of exons within its C-terminal myosin-tail homology domain (Datta et al., 2019). In contrast, the CEP290^KO^ model was made by insertion of a *β-Gal* cassette replacing exons 1-4 and causing complete loss of Cep290 protein production (Rachel et al., 2015). As previously reported, these two CEP290 mouse models displayed somewhat different retinal defects; therefore, we included both lines of mice in the present study. As described below, we found that both mouse models displayed photoreceptor degeneration and various disturbances in cilia function and OS formation. Several differences were observed between the two mutants, providing new insights into CEP290 functions in photoreceptor ciliogenesis, OS formation, CC trafficking, and Y-link/ciliary necklace stabilization.

## Results

### Immuno-Electron Microscopy reveals CEP290 localization between axoneme and membrane throughout the Connecting Cilium

We performed TEM imaging after immunogold staining of adult (P30) WT mouse retina with antibodies recognizing the Carboxyl-terminus (C-term) or Amino-terminus (N-term) of CEP290 (Fig. 1; for antibody validation, see Fig. S1), adapting an immunostaining protocol optimized for mouse rod CC antigens (Robichaux et al., 2019, Moye et al., 2023). For comparison, we also imaged retinas similarly stained with antibodies specific for another ciliopathy protein, RPGR. RPGR may help stabilize the Y-link complexes, is a proposed interactor of CEP290 in retina, and is also linked to ciliopathies and inherited retinal degenerations (Chang et al., 2006, McEwen et al., 2007, Anand and Khanna, 2012, Rachel et al., 2012, Sayer et al., 2006, Tsang et al., 2008, Megaw et al., 2015). Two isoforms of RPGR are expressed in the retina, a constitutive form (containing 19 exons; RPGR) and a retina-enriched isoform (stopping at exon 15 but containing a large portion of intron 15; RPGR^ret^) (Kirschner et al., 1999, Hong and Li, 2002). We used an antibody that recognizes only the retina-enriched isoform. Nanogold secondary antibodies were silver enhanced for visualization and are hereafter referred to as silver enhanced gold cluster, SEGC.

**Figure 1.**
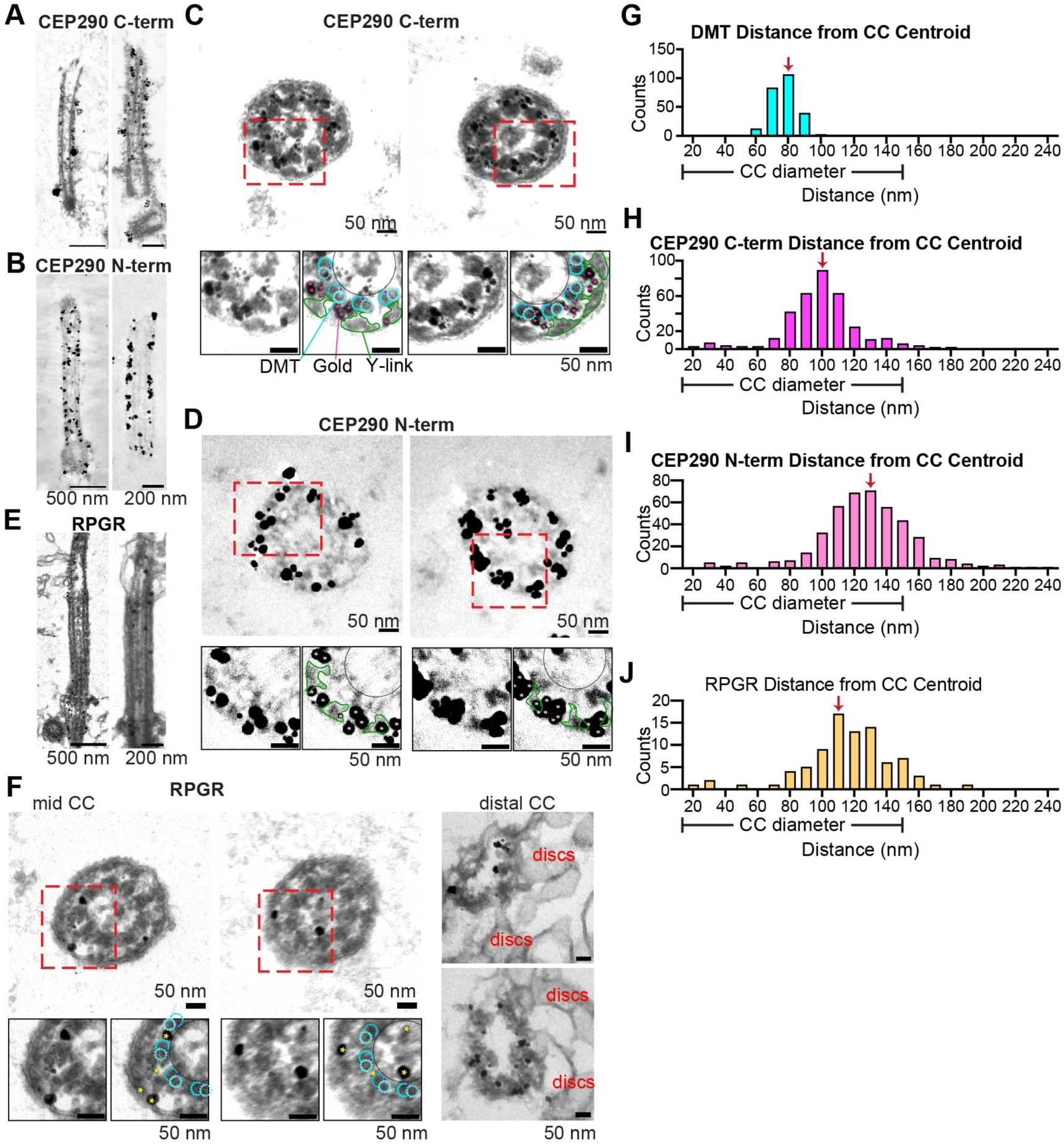
Localization of CEP290 and RPGR in WT mice using immuno-electron microscopy. (A-E) Immuno-EM of adult WT murine photoreceptor CC showing immunogold localization by antibodies specific for CEP290 C-terminus (A, C) or N-terminus (B, D), or by RPGR antibody (E, F). Longitudinal sections (A, B, E; scale bars = 500 nm in left panels and 200 nm in right panels) and perpendicular cross-sections (C, D, F; all scale bars = 50 nm) are shown. In the higher-magnification cross-sections, densities corresponding to microtubule doublets (cyan dots) and Y-links (green lines) that could be identified are indicated in overlays to the right, along with magenta stars indicating centers of SEGC. (F-I) Histograms representing radial distances of DMT and SEGCs from the centers of cilium cross-sections. (DMT measurements from n=27 cross sections from 5 animals, CEP290c from n=15 cross sections from 2 animals, 352 SEGCs; CEP290n from n=20 cross sections from 2 animals, 420 SEGCs; RPGR from n=14 cross sections from 1 animal, 85 SEGCs).

Sections roughly parallel to the axis of the CC displayed SEGCs near the ciliary membrane, with longitudinal distribution fairly uniform throughout the CC for CEP290 (Fig. 1A, B). RPGR (Fig. 1C, F) displayed a less robust staining than for CEP290, in that most transverse sections displayed only 2 or 3 SEGCs, but staining extended further up into the distal CC, where OS discs start to form, compared to CEP290 staining. Though staining of RPGR in the BB was also observed, this BB staining was also present in the RPGR^KO^ mice (Megaw et al., 2024). From multiple micrographs of cross-sectional views (Fig. 1C, D, K), we assembled histograms (Fig. 1G-J, Fig. S2) of radial positions of SEGCs, relative to the geometric centers of imaged cilia. Radial distributions of each SEGC, or of the DMTs, from the centers of each CC were calculated for all the transverse gold-labeled images that were near-circular (elliptical sections were not included, Supplemental Figure S2A). In contrast to the doublet microtubules (DMT) of the axoneme, which had a tight radial distribution centered at 80 nm, the CEP290 C-terminal antibody yielded a broader distribution, centered at 100 nm, and the distribution of the CEP290 N-terminal antibody, while similarly broad, was centered at 130 nm. The observation of smaller radial distances between the C-terminal antibody and the CC centroid, compared to the N-terminal antibody, was consistent with previous data showing that the C-terminus of CEP290 interacts with microtubules while the N-terminus interacts with the ciliary membrane (Drivas and Bennett, 2014). The distribution for RPGR was similarly broad with a center between 110 and 120 nm.

To complement the localization information from immunogold staining we used immunofluorescence microscopy in enhanced resolution modes (Fig. 2). Expansion microscopy (Robichaux et al., 2019, Moye et al., 2023) and Stochastic Optical Reconstruction Microscopy (STORM (Robichaux et al., 2019, Moye et al., 2023)), of adult mouse rods revealed individual DMT and CEP290 close to the DMT along the length of the CC (Fig. 2A). STORM imaging confirmed a radial distribution of CEP290 beyond the axoneme lumen marker, centrin, for both CEP290 antibodies (Fig. 2B) and RPGR (Fig. 2C). Iterative expansion microscopy (iUExM) of a human retina sample confirmed discrete CEP290 (Fig. 2 D, F and Movie 1) and RPGR (Fig. 2E, G and Movie 2) puncta along the lengths of the DMT in human rods (Fig. 2D, E) and cones (Fig. 2F, G). Cross-sectional views displayed close association of both antigens with the DMT (Fig. 2H, I).

**Figure 2.**
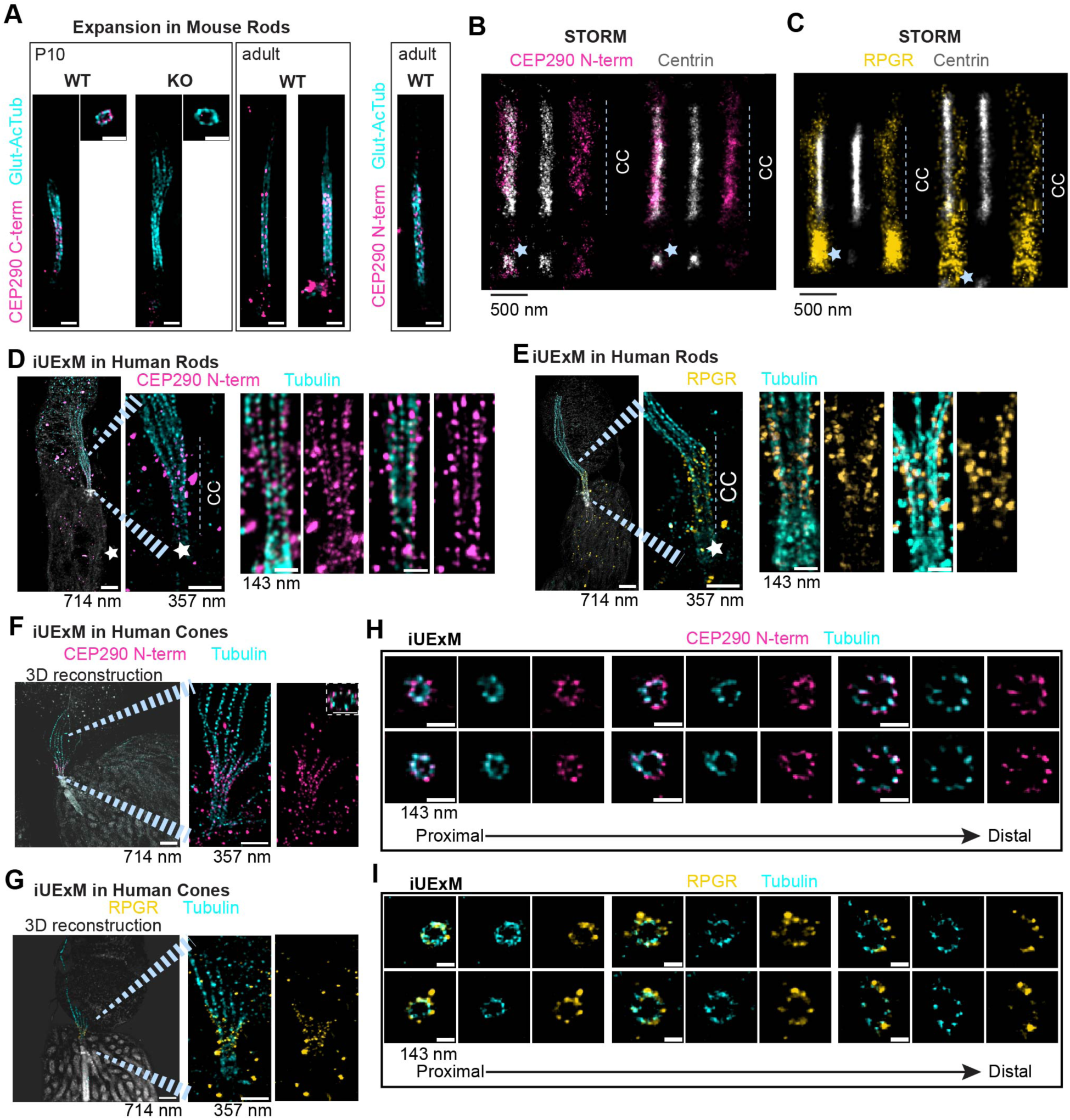
Superresolution Fluorescence imaging of CEP290 and RPGR in murine and human retina. (A) Expansion microscopy SIM images of mouse retina stained for CEP290 (C-terminus or N-terminus as indicated), in WT and CEP290^KO^ at P10, and in WT adults. Scale bar = 250 nm (corrected for 4x expansion). (B, C) STORM images of cilia (computationally straightened) stained for Centrin (grey) and either CEP290 N-terminus (B, magenta) or RPGR (C, gold) in WT adult mouse retina. Star = Basal Bodies. (D-J) are iterative expansion microscopy (iUExM) confocal images from expanded human retina of photoreceptor cilia immunostained for tubulin (cyan) and CEP290 N-terminus (D, F, magenta) or RPGR (E, G, gold). Lower magnification panels (left), DAPI (grey) is used to display non-specific membrane and rootlet staining. Scale bars (corrected for ∼14x expansion) = 714 nm in low-magnification panels, 357 nm in zoomed-in panels, and 143 nm for far-right higher-magnification examples in D and E. In H and I are cross-sectional slices through different regions of the cilia, corresponding to Proximal CC (PCC), Connecting Cilium (CC), or Distal CC (DCC). Scale bars = 143 nm (corrected).

### Ciliary necklace beads and ridges lost in CEP290^KO^ Connecting Cilia

Because of the proximity of CEP290 to the Y-links and to the DMT, which the Y-links connect to the membrane, we examined closely the Y-links and the ciliary membrane protrusions (ridges) associated with them that form the ciliary necklace (Zhang et al., 2023) in WT and CEP290 mutants. The ciliary necklace, a feature characterized from scanning EM and freeze-fracture on the membrane of primary cilia TZ and photoreceptor cilia CC that appears to look like a string of beads (Ringo, 1967), is hypothesized to be a part of the Y-link structures, as a transmembrane protein complex (Wensel et al., 2021, Zhang et al., 2023). We imaged mouse retinas by TEM at P10, before severe onset of the rapid retinal degeneration in CEP290 mutants, but after the CC are formed in WT. In reporting the results, we refer to three regions of the CC: Proximal CC (PCC) is defined as the 200 nm region from the last incomplete triplet to the end of the ciliary pocket where Y-link spacing changes ((Zhang et al., 2024), the mid-CC as the 900 nm region from the PCC to the bottom of the MT bulge, and the Distal CC (DCC) is defined as the 400 nm region from the start of the MT bulge to the end of the nascent discs.

These regions were identified in cross-sectional view by specific features associated with each region. The PCC had one side open to the IS cytoplasm, possibly displaying BB triplet MTs and distal appendages, and the other side contained nearly symmetric 9-DMTs with Y-links attaching to a distinct ciliary membrane. The mid-CC was identified by symmetrical 9-fold DMT assembly, with Y-links and an inner scaffold ring, fully encompassed by a distinct ciliary membrane. The DCC was identified as somewhat circular, with remnants of Y-links on some microtubules, possibly being connected to OS discs on one side, if present, and loss of the inner scaffold ring.

As reported previously, the Y-links were often present at all ages at which distinct ciliary membranes could be identified in both WT and *Cep290* mutants (Fig. 3A). Even upon detergent extraction, which was used to remove the lipids of the CC and thereby enhance visualization of the Y-links, there were robust (presumably protein-based) Y-link structures remaining in the Mid CC of both WT and CEP290^KO^ photoreceptors (Fig. 3B). However, the Y-links in the CEP290^KO^ cilia were much fewer and frequently of altered morphology, *e.g.*, fewer associated extracellular “beads”. The morphological distortions and “bead” number appeared to vary along the ciliary axis (proximal to distal), but this variation was difficult to quantify given the lack of precision in longitudinal localization of each section imaged.

**Figure 3.**
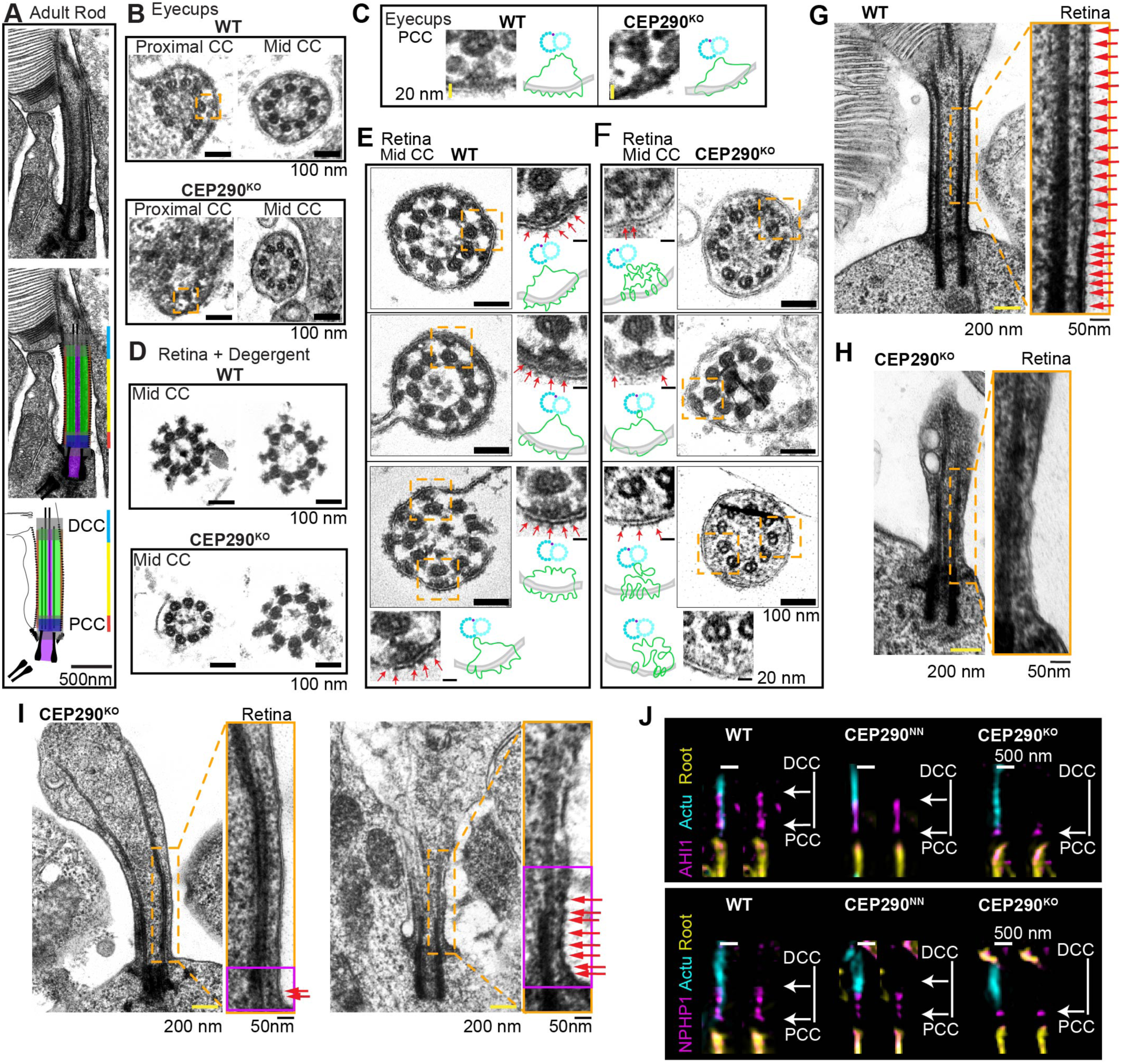
Effects of CEP290 deficiency on Y-links and ciliary necklace in the connecting cilium. (A-E) TEM images of cross-sections of CC in WT and CEP290^KO^ photoreceptors. (A) P10 eyecups from WT and CEP290^KO^, sectioned in the proximal CC or mid CC, with MTD and “Y-links.” The orange box highlights the region used for panel (E). (B) Images from P10 WT and CEP290^KO^ retinas incubated with Triton X-100 detergent to strip away the membranes, revealing the detergent-resistant components of “Y-links;” scale bars = 100 nm. (C, D) TEM images of mid-CC from P10 retina without detergent extraction. Higher magnifications display DMTs with attached Y-links, with ciliary bead ridges marked with arrows. Note the differences in the ciliary bead “ridges” between WT and CEP290^KO^ CC. (E) Higher magnifications from (A) highlighting similar appearance of proximal CC Y-links from WT and CEP290^KO^. Scale bars = 100 nm, zoomed insets = 20 nm. (F-H) TEM images of longitudinal sections of P10 eyecups from WT (F) and CEP290^KO^ (G, H) retina. Almost all WT CC membranes display ciliary necklace beads (arrows) throughout the CC, whereas CEP290^KO^ CC display beads in the proximal CC only, and often none are visible (*e.g.*, G). Scale bars: low mag = 200 nm, high mag = 50 nm. (I) Representative fluorescence (SIM, with computational straightening) images of cilia from WT, CEP290^NN^, or CEP290^KO^ retina showing localization of transition zone proteins, AHI1 and NPHP1, throughout CC in WT and CEP290^NN^, in contrast to their confinement to the proximal CC in CEP290^KO^. Scale bars = 500 nm.

In addition to the eyecup staining and sectioning used for Figure 3A and 3E, we also prepared sections using retinas isolated from RPE of WT and CEP290 mutant mice (Figure 3B-D, 3F-H) at P10. WT mid-CC cross-sections showed distinct Y-links connecting the DMTs to the ciliary membrane, and the presence of ciliary bead ridges (as was described in (Zhang et al., 2023)) (Fig. 3C, red arrows). In CEP290^KO^, Y-link morphology was altered, in that the portion attached to the DMT was usually present, but the connections between the DMTs and membrane were not as distinctive and appeared to lack dense associations to the ciliary membrane. In addition, there was a reduction in size and number of extracellular membrane ridges corresponding to the ciliary necklace (red arrows in Fig. 3G). In general, knockout cross-sections had fewer and shorter ridges/necklace beads, and the ciliary necklace (observed in longitudinal images) was often absent or present only in the most proximal region of the CC, where we had reported previously (Zhang et al., 2024) that the spacing of the ciliary necklace is different from that in more distal regions of the ciliary membrane (Fig. 3H). In contrast, in TEM of WT (Fig. 3F) and CEP290^NN^ (Fig. S3A) sectioned longitudinally, ciliary necklace bead protrusions were observed along the entire length of the CC. These structural aberrations support the hypothesis that CEP290 serves to stabilize the connection between the DMTs and the mid-to-distal ciliary membrane, with possibly a less important role in the proximal CC.

To test the hypothesis that localization of other transition zone proteins depends on CEP290, we checked the localization of two transition zone proteins whose genetic deficiencies, like those in *CEP290*, are associated with Joubert syndrome, AHI1 and NPHP1 (Brooks et al., 2018, Cheng et al., 2012, Gana et al., 2022, Wang et al., 2018). Superresolution fluorescence (SIM) revealed that, indeed, in CEP290^KO^ CC, these proteins are restricted to the PCC, whereas in WT and CEP290^NN^ CC, they are distributed throughout the length of the CC (Fig. 3J). In contrast to previous reports that an AHI1 mutation does not affect ciliary localization of CEP290 (Lessieur et al., 2017, Cheng et al., 2012), our results indicate that proper AHI1 localization in the CC depends on CEP290. The NPHP1 results are consistent with a report of genetic interactions between *Cep290* and *Nphp1* (Datta et al., 2021). We examined another TZ protein, NPHP8/RPGRIP1L, which, dissimilarly to AHI1 and NPHP1, localizes only to the PCC in WT photoreceptors (Arts et al., 2007), and found that the localization was unaffected in the CEP290 mutants (Fig. S4A), further supporting the idea that the PCC is less perturbed than are more distal CC regions by deficiencies in CEP290.

We also looked at the effects of the *Cep290* mutations on the distribution of RPGR, as it may also help stabilize the Y-link complexes and is a proposed interactor of CEP290 (Chang et al., 2006, McEwen et al., 2007, Anand and Khanna, 2012, Rachel et al., 2012, Sayer et al., 2006, Tsang et al., 2008, Megaw et al., 2015). SIM imaging revealed that the retina-specific splice variant, RPGR^ret^, is mislocalized in CEP290 mutant CC, being largely absent from the proximal region near the ciliary rootlet in CEP290^NN^ (Fig. S4B) but being either absent or confined to a shorter proximal region in the CEP290^KO^. Immunoblotting verified that RPGR^ret^ protein is present in CEP290 mutant retinas. Additionally, RPGR^ret^ is polyglutamylated (Sun et al., 2016), so probing with the polyglutamate-specific antibody GT335 revealed that the glutamylated form of RPGR^ret^ was also expressed in CEP290 mutant retinas (Fig. S4C). These results indicate that uniform RPGR localization throughout the length of the CC requires the presence of full-length CEP290.

### Stalled Ciliogenesis in CEP290^KO^Rods

At P10, ciliary structures have formed in both WT and CEP290^KO^ retinas, although in the knockout, they are much fewer in number and consistently lack the attached OS disc membranes that are seen in WT (Fig. 3, Fig. 4). To interrogate the role of CEP290 during development of the rod sensory cilium, we used TEM to score the various ciliogenesis stages at earlier timepoints, P3 - when ciliogenesis begins in mouse photoreceptors (Salinas et al., 2017, Sedmak and Wolfrum, 2011), and P7 - when OS disc formation begins (Fig. 4A-C). In WT retinas at P3, the majority of ciliogenesis stages observed were either migrating centrioles/BBs or cilia that have just broken the surface of the plasma membrane (extruding cilia “EC”) (Fig. 4A). However, in CEP290^KO^ rods, the majority of ciliogenesis stages observed were migrating BB, with a significantly larger number of BB with a ciliary vesicle still attached (“BCV”), and significantly different proportions of ciliogenesis stages in the knockout as compared to P3 WT rods (Fig. 4B, C). Photoreceptor development is somewhat asynchronous, so not every cell has a BB with a CV at the same time, as observed previously (Sedmak and Wolfrum, 2011). Some of the BB that were identified could actually belong to one of the other categories identified, with the associated features not captured in the ultrathin section; this technical limitation should affect both WT and KO samples to the same degree.

**Figure 4.**
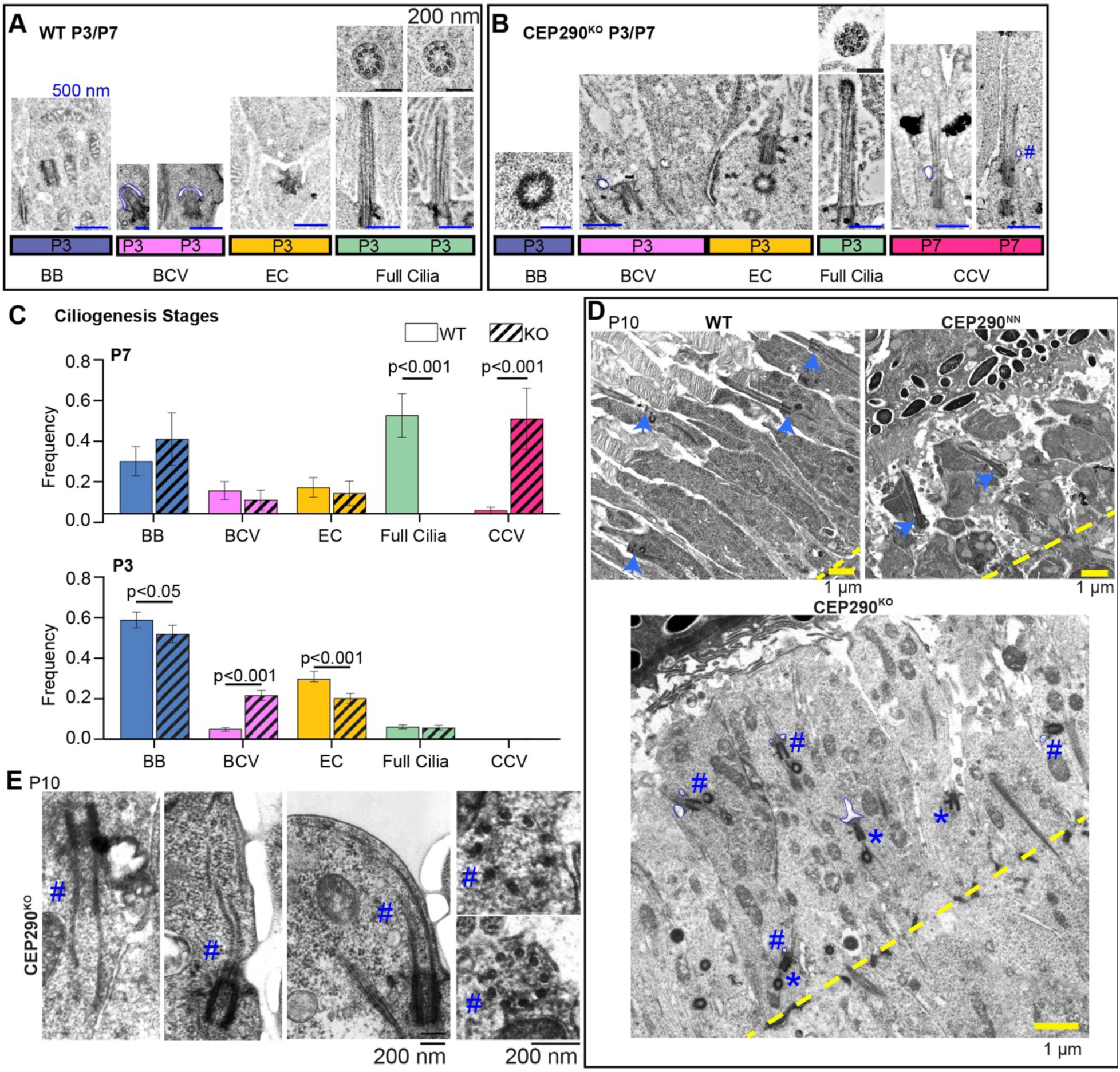
Impaired ciliary membrane formation in CEP290^KO^. (A, B) TEM images of P3 and P7 WT (A) or CEP290^KO^ (B) retinas showing different stages of ciliogenesis: “BB” = basal body only; “BCV” = basal body + ciliary vesicle; “EC” = cilia protruding from the inner segment; “Full cilia” = fully-extended CC with ciliary membrane; “CCV” = microtubules that have extended but still retain a ciliary vesicle. Scale bars: 500 nm (longitudinal sections), 200 nm (cross sections). (C) Plots of numbers of ciliogenesis stages observed at P3 and P7 in WT and CEP290^KO^ as frequencies ± s.e.m. WT P3: 370 BB, 187 EC,32 BB+CV, 39 Full Cilia; KO P3: 225 BB, 94 BCV, 88 EC, 25 Full Cilia, 1 CCV; WT P7: 16 BB, 7 BCV, 8 EC, 30 full cilia, 1 CCV; KO P7: 11 BB, 2 BCV, 3 EC, 14 CCV. For P3 and P7 only 1 animal was examined. T-tests were performed to compare each category; if no p-value is listed it was non-significant. (D) P10 retinas (n=4 animals for all genotypes, showing similar results) from WT, CEP290^NN^ CEP290^KO^ retinas. Arrows: developing OS with BB/CC located at the distal edge of the IS (WT and CEP290^NN^ only). Asterisks: BB with distal appendages located in proximal IS (CEP290^KO^ only); Pound sign: BB with distal appendages, axoneme and associated ciliary vesicles but no distinct ciliary membrane. Ciliary vesicles are outlined in blue in panel A, B, and D. Scale bars = 1 μm. (E) Higher magnification images from CEP290^KO^ at P10 of nascent cilia with extended axonemes but no distinct ciliary membrane. Two micrographs on far right are cross-sections.

At P7, when ciliogenesis is near completion and OS discs begin forming, about half of WT rods scored were “Full Cilia”, protruding from the inner segment (IS) with a complete ciliary membrane. In contrast, most ciliary structures identified in CEP290^KO^ rods were either BBs (no CV and not protruding from the IS), or cilia with a CV, “CCV” (full cilia that still possess a CV on at least one side of the axonemal microtubules) (Fig. 4A-C). At P10, we observed CCV in CEP290^KO^ rods, whereas in WT and CEP290^NN^ rods, only Full Cilia were observed (Fig. 4D). Upon closer investigation, it appeared that the CCVs found in CEP290^KO^ rod ISs at P7 and P10 had one side with a membrane and one side exposed to the IS cytoplasm (marked with # in Fig. 4D, E). There were also many examples at P10 of CEP290^KO^ “cilia” with a complete absence of ciliary membrane (just extended microtubules within the cytoplasm) (Fig. 4 E, the full micrographs are displayed in Fig. S5). Even at P10, there are striking differences in the stages of ciliogenesis observed in CEP290^KO^ *vs.* WT rods (Fig. 4D-F).

To complement our TEM analysis, we used fluorescence and structured illumination microscopy (SIM) to assess ciliary vesicle formation at P3 in CEP290^KO^ retinas. The number of fluorescent puncta that were positive for the distal appendage vesicle (DAV) markers, CP110 and CEP97 (which are also potential CEP290 interactors (Tsang et al., 2008, Kobayashi et al., 2014)), were not significantly different in CEP290^KO^ retinas compared to WT (Fig. S5B, C), indicating normal DAV protein recruitment and formation. Furthermore, a distal appendage protein, CEP89 (Yang et al., 2015), localizes normally in WT and CEP290 mutant photoreceptors to the PCC (Fig. S5D). These results indicate that loss of CEP290 stalls ciliogenesis after CV formation and before ciliary membrane formation, but CEP90 loss does not disrupt microtubule extension of the axoneme.

### Massive accumulation of extracellular vesicles in Cep290 mutants

To address the consequences of ciliogenesis defects at a developmental time when extensive outer segment disc formation has normally occurred (P10), retinal morphology from WT, CEP290^NN^, and CEP290^KO^ mice was examined by TEM (Fig. 5). In WT retinas, fully formed CC and the presence of OS discs were observed (Fig. 5A). WT OS disc formation begins at P7, with fully formed OS by P21 (LaVail, 1973). However, in the retinas from CEP290 mutants at P10, numerous extracellular vesicles (EVs) were observed instead of discs in the OS layer (Fig. 5A, B). In CEP290^NN^ retinas, some OS discs were found (Fig. 5B, yellow stars), but the EVs were generally more prevalent. The average diameter of the EVs in the CEP290^NN^ retinas was 200 nm ± 72.64 (Fig. 5C), which is similar to the mean diameter of the EVs detected in the *rds* mutant mouse (Molday and Goldberg, 2017, Salinas et al., 2017). CEP290^KO^ retinas also had EVs, but overall fewer than in the CEP290^NN^ retinas.

**Figure 5.**
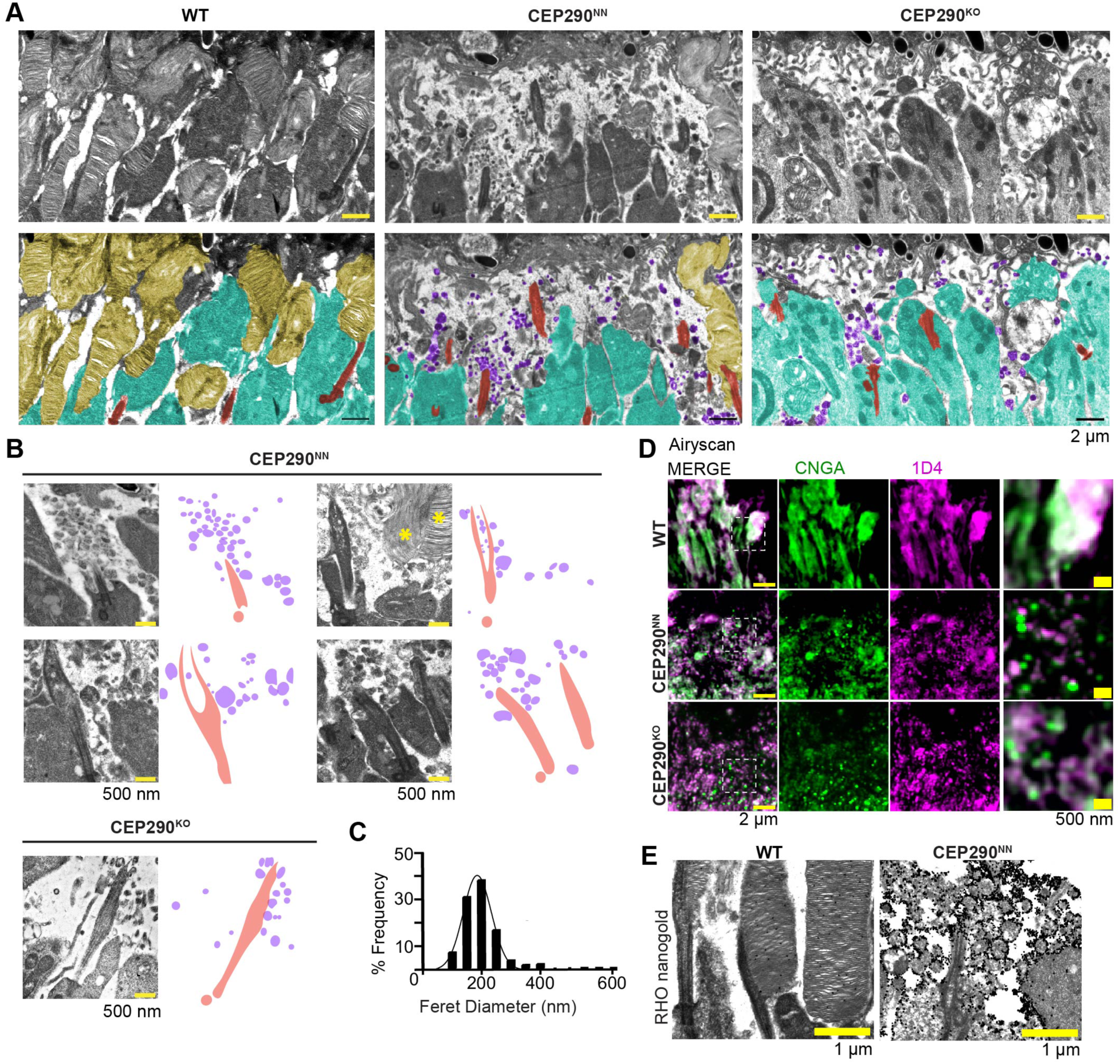
Extracellular vesicles (EVs) in CEP290 mutant mouse retina. (A) TEM images from wildtype (WT), CEP290^NN^, and CEP290^KO^ showing regions corresponding to the outer segment at P10, with greyscale (upper panels) and pseudo-colored (lower panels) representations: OS – yellow, IS – cyan, CC/BB – red, EVs – purple. Scale bar = 2 μm. (B) TEM images of EVs in mutant OS, with pseudo-colored visualization to the right. Scale bars = 500 nm. Asterisks indicate rarely observed objects resembling OS discs in CEP290^NN^. (C) Histogram with best-fit line to normal distribution of EV Feret diameters measured from CEP290^NN^. Mean = 200 nm +/- 72.46 (s.d.), n= 578 EVs from 2 CEP290^NN^ mice. D) Airyscan images of the photoreceptor layer of fixed retinal cryosections at P10 from WT, CEP290^NN^, and CEP290^KO^ retinas, immunostained for CNGA and rhodopsin (mAb 1D4); scale bar = 2 μm. Areas in white dashed boxes are shown at 3.5 x higher magnification at far right; and dimensions of far right panels = 3 μm. (E) Immunogold labeling of rhodopsin (1D4) in TEM images from WT and CEP290^NN^ retinas at P10. 1D4 labels OS discs in WT and EVs in CEP290^NN^. Experiment was replicated 3 times. Scale bars = 1 μm.

To determine potential protein content of these vesicles, we used immunofluorescence (IF) with Airyscan confocal microscopy of OS proteins in P10 WT and CEP290 mutant retinas. Based on our EV size determination from TEM, we looked for fluorescent puncta whose diameters were at or near the resolution limit of ∼150 nm (Fig. 5 D, showing only OS region). Very few, if any, such puncta were observed in WT, but there were multiple EVs visualized by IF in both CEP290^KO^ and CEP290^NN^ retinas. In both mutants, many of these puncta had strong signal for the rhodopsin antibody, 1D4, which could represent EVs or aggregates of rhodopsin mislocalized in the IS, as observed previously (Potter et al., 2021b). In fluorescence images at this resolution, staining of EVs cannot be unambiguously distinguished from rhodopsin mislocalized within the IS. Therefore, the presence of Rhodopsin in the EVs was confirmed through immunoelectron microscopy using the 1D4 antibody (Fig. 5E and Fig. S6A, B).

There was also strong punctate signal in both CEP290 mutants for the α subunit of the cyclic-nucleotide-gated channel, CNGA, a trans-membrane protein of the OS plasma membrane (Fig. 5D). Most puncta stained for both antigens, but a subset stained strongly for CNGA with weak, if any, 1D4 staining (zoomed-out images in Fig. S7A). In sections from the same CEP290^NN^ eyecups as were used for CNGA and 1D4, little evidence was observed for extracellular vesicle staining for the disc rim tetraspannin protein, peripherin (PRPH2) and for phosphodiesterase-6 (PDE6β); whereas in CEP290^KO^, some PRPH2 puncta were observed. What appears to be disc staining was observed for both PHPH2 and PDE6β in CEP290^NN^ retinas (see also Fig. S7B, C).

Immunofluorescence for cone markers demonstrated no significant decrease in the number of cones, with cone OSs appearing to be generally intact in both CEP290 mutants, based on peanut agglutinin (PNA) and cone arrestin (cArr) staining (Fig. S7D). There were no indications of EVs observed in any cone marker staining for either mutant. These results suggest a mechanism for EV formation in CEP290^NN^, possibly from initiation but inefficient completion of disc formation, that is missing in CEP290^KO^ in which no discs and few EVs form. It may be that CEP290 plays a less important role in OS formation in cones, although we did not identify cone OS in electron micrographs, likely due to the very low cone number, compared to rods, in mouse retina.

### Mislocalization of centrins, luminal scaffold components, but not of INPP5E, a ciliary membrane-associated antigen, in Cep290 mutants

Because we have observed distinct differences in the photoreceptor defects caused by truncation vs. loss of CEP290, we looked for differences in markers of the ciliary lumen and ciliary membrane, which is largely lacking in CEP290^KO^ (Fig. 4E). We performed immunofluorescence with SIM and STORM using antibodies that recognize 1) Inositol polyphosphate-5-phosphatase E (INPP5E), a ciliopathy-related protein known to localize to the CC membrane surrounding the axoneme (Sharif et al., 2021) as well as to membranes within the IS (Fig. 6A and Fig. S8A-C); and 2) multiple isoforms of centrin, a calcium-binding protein that generally serves as a marker for the BB in primary cilia but which localizes throughout the lumen of the CC in photoreceptors (Robichaux et al., 2019), and which is unlikely to form constitutive complexes with CEP290.

**Figure 6.**
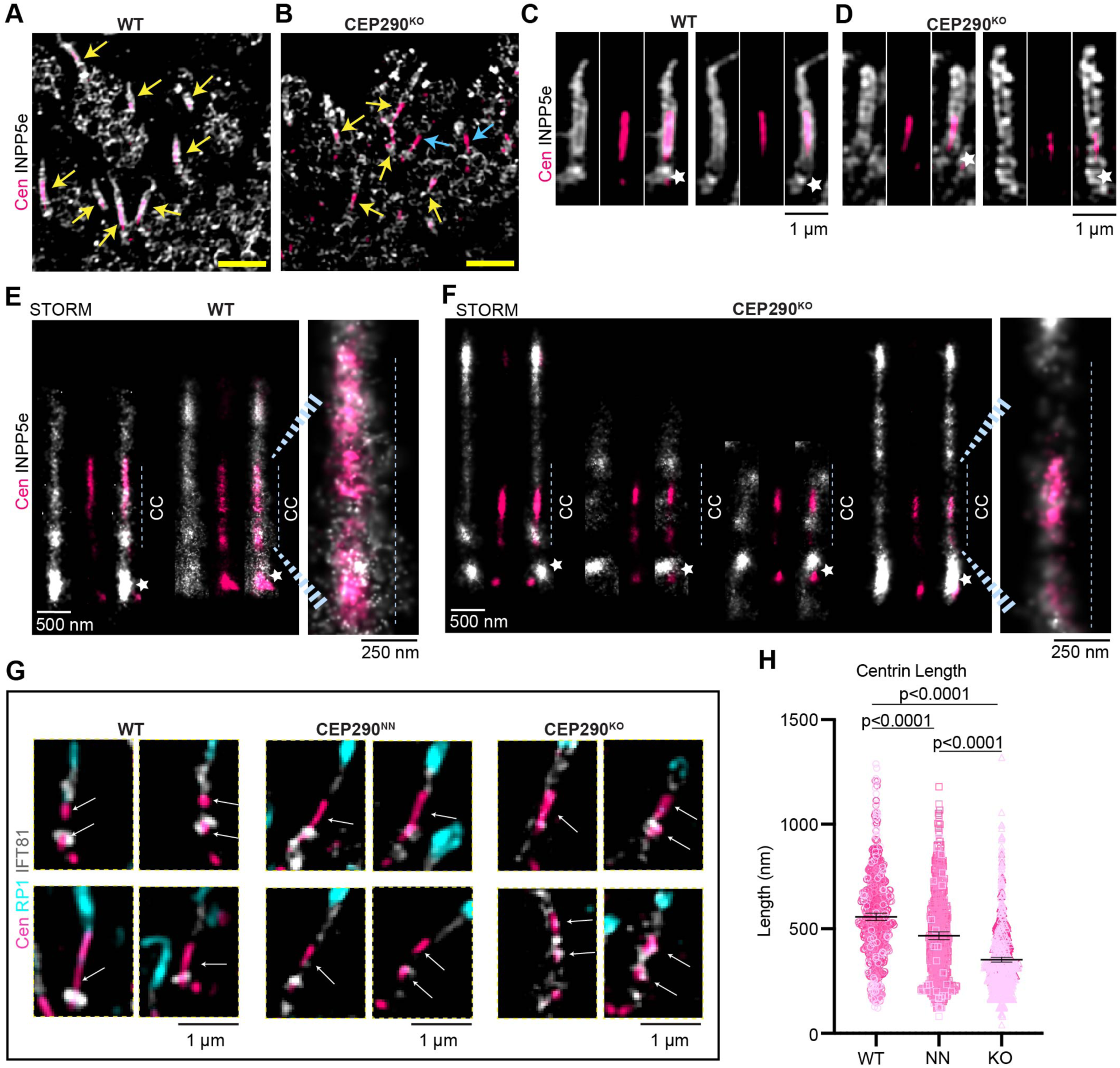
Effects of CEP290 deficiency on spatial distributions of CC proteins. (A, B) SIM images of retinas immunostained for phosphoinositide phosphatase, INPP5E (gray), and centrins (identified with pan-centrin antibody, pink) from mice of indicated genotypes. Yellow arrows point to INPP5E staining CC and axonemes, blue arrows indicate centrin staining with no INPP5e surrounding. Scale bars = 2 𝜇m. SIM (C-D) and STORM (E-F) images of straightened, individual cilia, highlighting INPP5e localization in the CC, and centrin abnormalities in the CEP290^KO^ photoreceptors. Each SIM image in C and D is 1 𝜇m wide, and each STORM image in E and F is 500 nm wide. Star = basal bodies. (G) SIM images displaying centrin labeling in the context of ciliary markers RP1 (cyan) and IFT88 (gray) in WT and CEP290 mutant retinas, showing varied distribution of centrin labelling at P10. (H) Scatter plot displaying lengths of centrin from individual cilia in SIM images, displaying 95% CI, and p values from *T*-test showing pairwise significance of differences in centrin length between genotypes. WT n = 609, NN = 545, KO = 864 cilia, all from either 3 or 4 different mice. Mean values: WT = 556.6 nm, NN = 466.5 nm, KO = 351.9 nm.

The localization pattern of INPP5E staining in the CEP290 mutants was surprisingly similar to that in WT. Interestingly, INPP5E staining appeared to extend into the axoneme where nascent OS discs are formed (in WT), with similar length of staining detected in the CEP290^KO^. This staining could represent the possibly less affected cone cilia, or it could represent cases of FULL cilia observed in the CEP290^KO^ (for example, as seen in the left TEM from Fig. 7B). Thus, at least some of the properties of the CC membrane are intact in the CEP290^KO^ when emerging from the IS, whereas other features, such as the ciliary necklace, are missing. In addition, centrin staining that was not surrounded with INPP5E (Fig. 6B, blue arrows) was observed, which could represent BB staining or axonemes forming within the IS without associated membrane.

**Figure 7.**
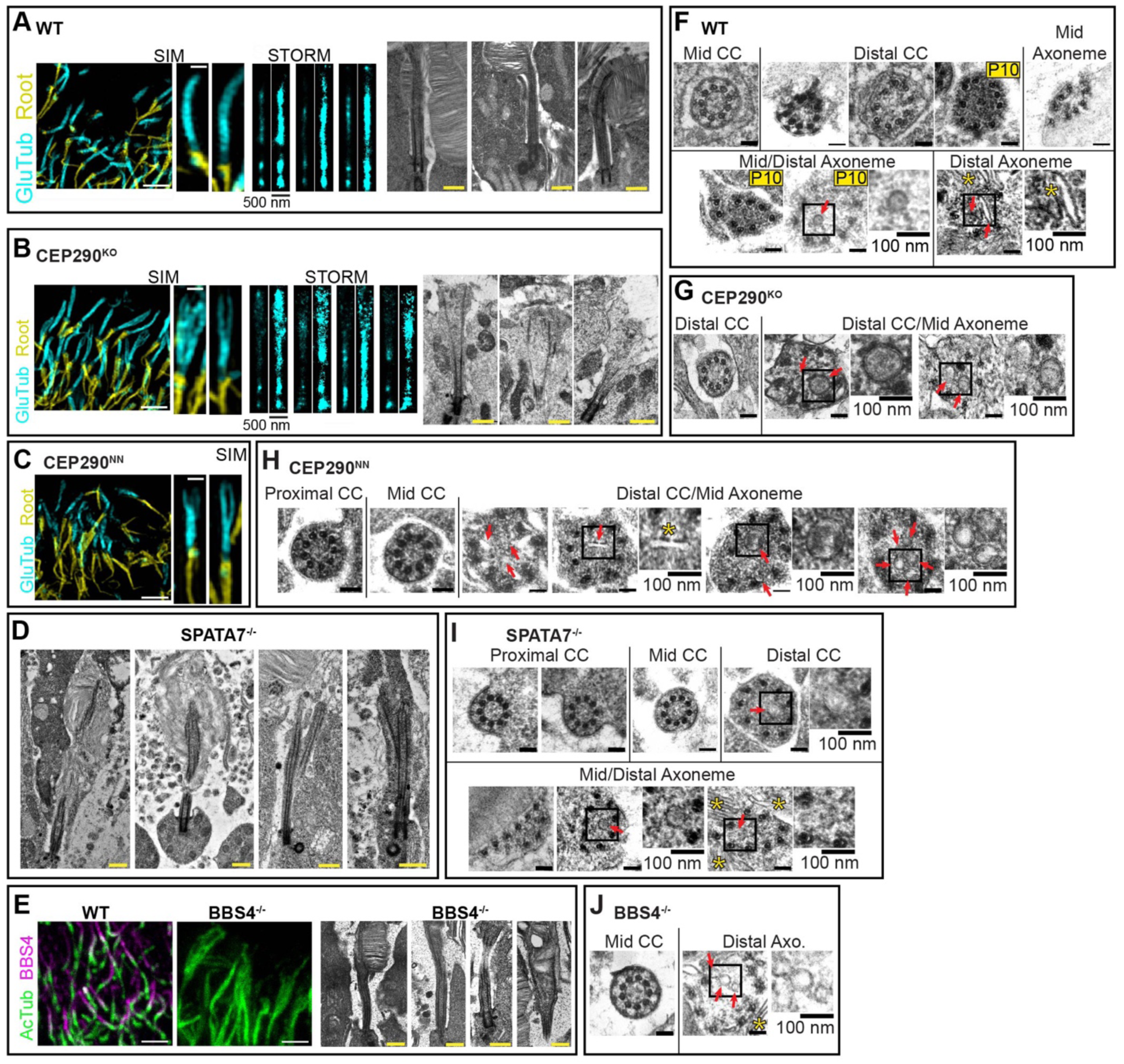
Ciliary axoneme splaying in ciliary mutant photoreceptors. (A, B) SIM (left), STORM (middle), and TEM (right) images of cilia from WT (A) and CEP290^KO^ (B) retina at P10. SIM images display staining of glutamylated tubulin (GluTub) and rootletin. STORM images display staining of glutamylated tubulin, with overexposed versions to the right. Scale bar = 2𝜇m (low mag SIM), straightened SIM and STORM cilia are all 500 nm wide, and TEM = 2 𝜇m. (C) Confocal from WT and BBS4^-/-^ retinal cryosections at P30. To the right are TEM micrographs from BBS4^-/-^ retinas at P30. Scale bars = 2 𝜇m. (D) TEM micrographs from SPATA7^-/-^ retina at P10. Scale bars = 500 nm. (E-H) Electron micrographs showing transverse sections through the CC and axoneme of photoreceptors in WT (adult and P10), and different ciliary mutants (F-H). Arrows and insets highlight membranous features found in ciliary lumen, with disc-like structures indicated with yellow stars. Scale bars = 100 nm.

Differences were observed in centrin localization in both CEP290 mutant retinas at P10, but more severely in CEP290^KO^. Specifically, there were cilia detected (through INPPE staining) wherein there was no centrin labelling, and often centrin staining appeared shorter or fragmented compared to WT (Fig. 6A-G). Of note, centrin staining length can vary at P10, even in WT, as these are not fully mature photoreceptors, therefore, fragmented (*i.e.*, centrin labeling DCC and PCC discontinuously in the same cilium), and short centrin labeling was also observed in WT at P10 (Fig. 6G, with zoom-out images in Fig. SD-F), but to a much lesser extent than in the mutants. Measurements of centrin length from SIM images showed that there was a significant difference between WT and CEP290^KO^, as well as in the CEP290^NN^ retinas, compared to WT (Fig. 6H), further indicating that in CEP290^KO^ retinas, photoreceptor ciliogenesis is stalled.

### Microtubule splaying in photoreceptor axonemes of CEP290 mutant mouse rods

There is an inner scaffold (Mercey et al., 2022, Zhang et al., 2024) within the CC axoneme where centrins reside, that has been proposed to “zip” the microtubule doublets together and help maintain axonemal symmetry, geometry, and stability (Le Guennec et al., 2020, Mercey et al., 2022). Microtubule splaying in photoreceptor axonemes has been described in multiple ciliopathy mouse models, and is often coupled with ciliary protein mislocalization, instability of the axoneme, and aberrant OS disc morphogenesis (Dharmat et al., 2018, Mercey et al., 2022, Faber et al., 2023). Super resolution methods (SIM and STORM) and TEM were used to examine the ultrastructure of rod axonemes in WT and CEP290^KO^, as well as other ciliopathy mutant mouse retinas. For SIM and STORM, immunolabeling with anti-GT335, an antibody that binds to polyglutamylate, and therefore the polyglutamylated MTs in the CC, was used to visualize photoreceptor axonemes in WT and CEP290 mutant rods at P10 (Fig. 7A-C). MT splaying was evident in many CEP290^NN^ and CEP290^KO^ photoreceptors, as demonstrated by SIM, STORM, and in electron micrographs (Fig. 7B, C).

For comparison, S*pata7*^-/-^ retinas were also examined. SPATA7 is a ciliopathy-associated protein which localizes throughout the length of the CC and for which centrins and other CC antigens also displayed localization defects (∼750 nm length of centrin staining in S*pata7*^-/-^, compared to ∼1200 nm in WT STORM images at P15 (Dharmat et al., 2018)). Indeed, at P10, which precedes major photoreceptor degeneration, MT splaying was observed (Fig. 7 D). We had previously observed MT splaying in *Spata7*^-/-^ by cryo-electron tomography at P15 (Dharmat et al., 2018), but this is its first observation by conventional TEM.

To examine the possibility of MT splaying in a ciliopathy mutant generated from loss of a trafficking protein, rather than more structural ones, we examined *Bbs4^-/-^* retinas. BBS4 localizes to the CC of rod photoreceptor cells (Zhang et al., 2014). Confocal microscopy of retinal cryosections immunolabeled with an antibody targeting acetylated tubulin (AcTub, a marker for CC microtubules) and TEM at P30 (precedes major photoreceptor degeneration (Mykytyn et al., 2004)) revealed MT splaying in the *Bbs4^-/-^*photoreceptors (Fig. 7E).

The cross-sectional ultrastructure of photoreceptor axonemes from mid-CC to distal axoneme from WT and the cilia mutants mentioned above were also analyzed in electron micrographs to visualize the splaying from a transverse view (Fig. 7F-J), using the definitions of the PCC, mid-CC, and DCC as described for Fig. 5. Additionally, in classifying these sections according to axial position, the OS axoneme was identified by microtubules within and adjacent to OS discs that have a non-symmetrical, often triangular shape, and consisting of some singlet microtubules but mostly doublets, and no presence of Y-links or inner scaffold ring. In those classified as distal axoneme there were not always 9 microtubules present; mostly singlets and a few doublets were observed, sometimes associated with unknown membranous structures in the OS (red arrows, Fig. 7F-J). CC ultrastructure was compared in adults as well as at P10.Although OS discs are still forming and the axoneme is actively extending at P10, P10 WT CC ultrastructure was comparable to that in adult WT. In *Cep290*^NN^ and *Cep290*^KO^ transverse cilium sections at P10, it was difficult to differentiate between distal CC and mid axoneme, due to the MT splaying, so they were grouped together. Many circular/disc-like membranous structures within the axonemal lumen (∼90% of 200 cross-sectional TEM images displayed these) were observed in the mid- and distal OS axoneme in both *Cep290* mutants (Fig. 7G, H). In all the WT images analyzed, only two instances of these membranous structures were observed, once in adult and once in P10 retinas, out of 200 TEM images (Fig. 7F). These membranous structures were similar to what was observed in longitudinal sections of *Cep290*^KO^, as seen in Fig. 7B.

Transverse CC axoneme ultrastructure was also analyzed by TEM in *Spata7*^-/-^ and *Bbs4*^-/-^ models. In *Spata7*^-/-^ retinas, at P10 and P15, the ciliary ultrastructure up to mid-CC was similar to WT (as was observed in longitudinal sections previously, (Dharmat et al., 2018)) (Fig. 7I). However, in the DCC and OS axoneme, where all 9 DMTs were still present, the membranous structures within the microtubule lumen were often present (∼75%, out of 30 images, 23 displayed axonemes with membranous structures), which was the same localization for these abnormal structures in P10 *Cep290* mutants. Abnormal membranous structures were also seen in *Bbs4*^--/-^ CC at 3 months (out of 5 transverse TEM images, 2 displayed them) (Fig. 7J).

MT splaying and luminal scaffold disruption within the CC, which is the trafficking highway of the photoreceptors, prompted investigation into intraflagellar transport (IFT) in CEP290 mutant photoreceptors. SIM was used to analyze the localization of three IFTs – IFT-A complex protein IFT140 and IFT-B complex proteins IFT88 and IFT81. IFT localization in adult CC is typically observed as 2 puncta, one at the PCC and one at the DCC, where there is hypothesized to be a second IFT “docking” zone, at which the velocity of IFT trains decreases (Yang et al., 2019, Oswald et al., 2018, Jensen et al., 2015, Nachury and Mick, 2019, De-Castro et al., 2022) for OS disc formation. However, at P10 in WT, we often observed higher intensity of the PCC IFT punctum and a fainter intensity of the DCC IFT punctum, likely due to the build-up at the CC base of cargoes required to be trafficked into the developing axoneme at this age. Differences in IFT protein localization were found between P10 WT and the CEP290 mutants. In WT photoreceptors, the IFTs almost exclusively colocalized with the ciliary marker, whereas in both CEP290^KO^ and CEP290^NN^ rods there were both punctate accumulations along the CC and axoneme, and smaller punctate dots outside of the ciliary ROI, possibly corresponding to EVs, for all three IFTs in both mutants (Fig. 8A-C).

**Figure 8.**
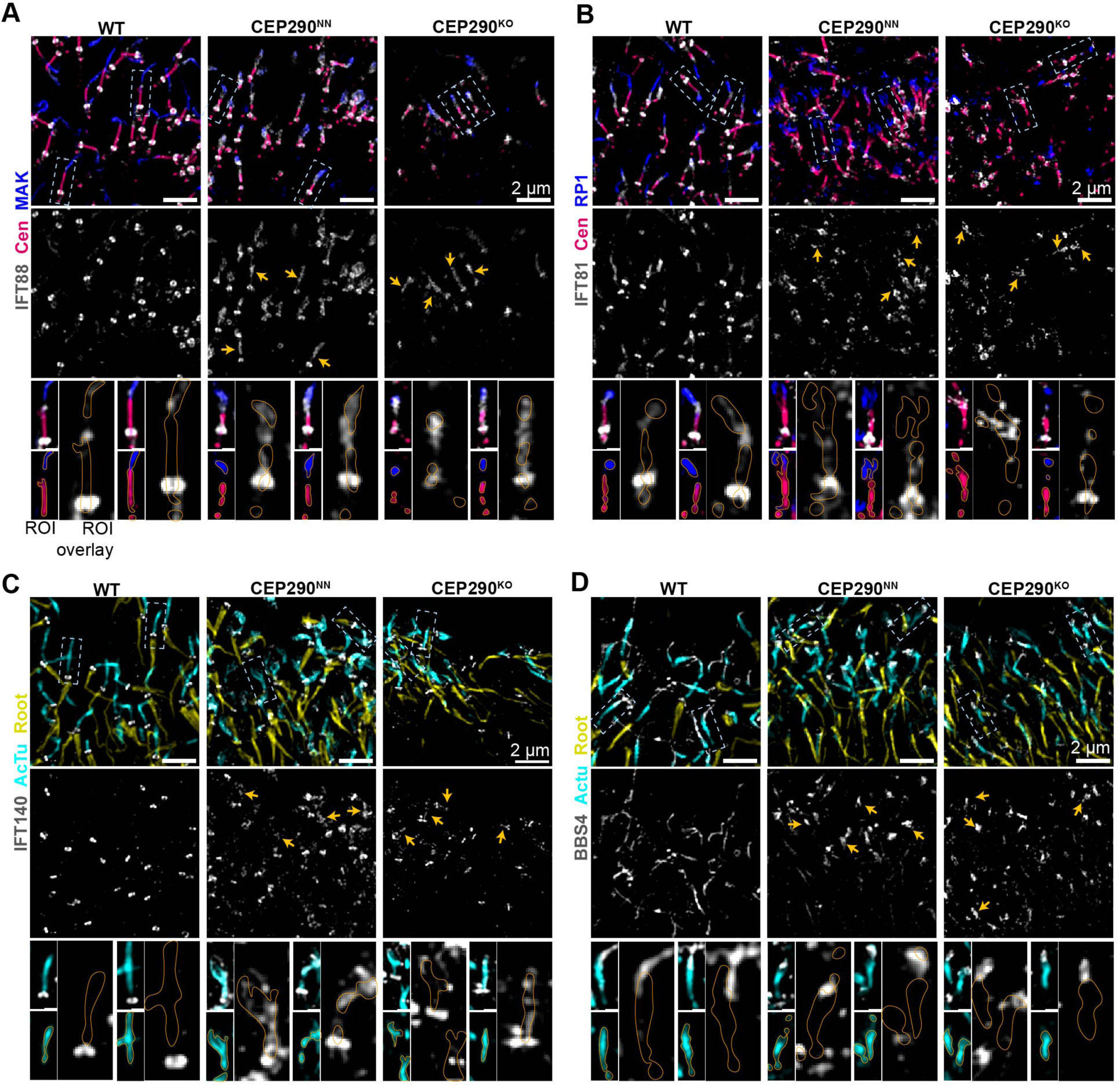
Accumulations of Intraflagellar Transport Proteins in the CC of CEP290 mutant retina. (A-D) SIM low magnification images of cilia from WT and CEP290 mutant retinal cryosections at P10, displaying localization of indicated ciliary proteins. Beneath each merged image is the IFT or BBS4 only. Scale bar = 2 𝜇m. White dashed boxes indicate the cilia chosen for zoomed panels, orange lines indicating the ROI of the ciliary staining. Each are 1 μm. Each staining was performed three separate times on sections from 3 or 4 different animals.

Conversely, the localization of BBS4, which is also involved in ciliary trafficking and localizes along the entire CC/axoneme in WT, appeared decreased in length in both CEP290 mutants (Fig. 8D). Shortened distributions in the mutants were also observed for retinitis pigmentosa 1 (RP1) and male germ-cell associated kinase (MAK), which localized to the photoreceptor ciliary axoneme (Omori et al., 2010, Liu et al., 2003, Liu et al., 2002, Moye et al., 2018) (Fig. 8A, B).

## Discussion

### CEP290 Photoreceptor Localization in Mice and Humans

In this study, CEP290 and RPGR were localized between the doublet microtubules (DMTs) and the ciliary membrane in the connecting cilia (CC) of both human and murine photoreceptors. Immunogold electron microscopy (EM) combined with fluorescence microscopy enabled precise protein localization relative to cellular structures and protein complexes. CEP290 immunolabeling revealed a symmetrical pattern near the Y-links in both species, consistent with previous immunogold EM findings of CEP290 in *Chlamydomonas* by Craige*, et. al*, in which most immunogold particles were seen in the region between adjacent Y-links (Craige et al., 2010), and findings from iterative expansion protocols (Louvel et al., 2023). Furthermore, our immunogold TEM results suggest N-terminal immunolabelling of CEP290 nearer the membrane, and C-terminal labeling nearer the MTs, aligning with *in vitro* observations using truncated constructs (Drivas et al., 2013).

### CEP290 and Y-links in the Connecting Cilium

CEP290 has been hypothesized as a structural scaffold of the Y-links in the TZ of cilia (reviewed in (Park and Leroux, 2022)). We previously found Y-link structures remain in the mid-CC of CEP290^NN^ and CEP290^KO^ photoreceptors, though with a shorter distance between DMT and membrane (Potter et al., 2021a), consistent with results for CEP290 loss in TZ of other ciliated cells (Craige et al., 2010). For the first time here, we observed disruptions in Y-link shape and connection to ciliary membrane throughout the mid to distal CC. Concurrently, we report a loss of ciliary beads along the length of the mid-distal CC and loss of ciliary bead globular ridges (transverse view) in the CEP290^KO^, structures that may be complexed with the Y-links. Mutations in RPGRIP1L and NPHP4 led to loss of Y-link connections and vesicle accumulation in the CC (Gogendeau et al., 2020), similar to CEP290^KO^ mutants. However, CEP290 mutants exhibited more profound disruptions in Y-link stability along the mid to distal CC in mice and humans. These results, along with the structural defects in mutant mice, support the hypothesis that CEP290 contributes to Y-link stability along the mid to distal CC.

### Role of CEP290 in MT Extension and Ciliary Membrane Formation in Photoreceptors

CEP290 is critical for photoreceptor ciliogenesis, as its ablation resulted in halted ciliogenesis and altered CC structures, in agreement with previous studies in RPE-1 cells or *Paramecium* (Kobayashi et al., 2014, Wiegering et al., 2021). In CEP290^KO^ photoreceptors, full-length axonemal structures which had altered ciliary membranes co-occurred with stunted axonemes, often captured with abnormally associated ciliary vesicles. While some photoreceptor CCs in CEP290^KO^ mutants contained inner scaffold rings with 9-fold DMT symmetry, membrane formation was frequently absent. Previous studies demonstrated similar phenomena of membrane-less axonemes or bulging lumens in patient-derived (LCA) organoids and (Joubert Syndrome) fibroblasts (Shimada et al., 2017). Surprisingly, the lipid phosphatase INPP5E, which was previously reported to display a decreased ciliary abundance in the absence of TZ proteins NPHP1 or RPGR in photoreceptors and kidney epithelial primary cilia (Rao et al., 2016, Ning et al., 2021), localized normally in CEP290^KO^ photoreceptors. The small GTPase, Rab8, has been shown to be required for the release of ciliary vesicles and subsequent ciliary membrane formation (reviewed in (Zhao et al., 2023, Chen et al., 2021, Tsang et al., 2008, Yoshimura et al., 2007, Nachury et al., 2007, Kim et al., 2008)). The unaffected localization of INPP5E in CEP290^KO^ photoreceptors and Rab8’s known interaction with CEP290 (Kim et al., 2008) supports the hypothesis that CEP290 is involved in ciliary vesicle-mediated ciliary membrane formation, but not strictly required for MT extension.

### Microtubule Splaying and Inner Scaffold Ring Integrity

CEP290 mutants displayed disrupted microtubule (MT) organization and inner scaffold ring stability. The inner scaffold ring has been shown to contain POC5, Centrin, FAM161A, and POC1, as revealed by immunolocalization (Mercey et al., 2022, Sala et al., 2024), and POC1b, as suggested by the phenotype of *Poc1b* mutants ((Beck et al., 2014, Durlu et al., 2014, Roosing et al., 2014, Patnaik et al., 2015)). Interestingly, how centrins, localized to the CC lumen, might interact with the inner scaffold ring is not fully understood, especially given that centrins are restricted to the BB in other primary cilia (Laporte et al., 2024) but TZ’s still possess an inner scaffold (Fisch and Dupuis-Williams, 2011). In CEP290^NN^ and CEP290^KO^ photoreceptors, the inner scaffold ring was not observed in the DCC, but remained unperturbed in the PCC and mid-CC. Additionally, instead of the uniform centrin staining observed throughout the CC, CEP290 mutant models revealed punctate distribution of centrins in the PCC and mid-CC. These findings align with similar phenotypes in other ciliary mutants, including those deficient in FAM161A and SPATA7, where disrupted centrin localization was reported (Dharmat et al., 2018, Mercey et al., 2022).

It has been hypothesized that one of the roles of the inner scaffold is to maintain the integrity, length, and circularity of the centrioles and CC (Le Guennec et al., 2020, Steib et al., 2020, Schweizer et al., 2021, Atorino et al., 2020, Sala et al., 2024). However, DMT splaying in photoreceptor CC is provoked by the ablation of multiple different ciliary proteins, including centrins (Fig. 7 and (Ying et al., 2019)), indicating that splaying could be a general sign of CC instability. It is unclear at this point if inner scaffold ring disruption and centrin mislocalization are related to DMT splaying, or if they occur simultaneously due to a different cause.

### Role in Extracellular Vesicle Formation

The presence of EVs and absence (KO) or perturbation (NN) of discs in CEP290 mutants supports the hypothesis that CEP290 plays a role in packaging membrane cargo into discs. EVs are observed in other mouse models displaying photoreceptor degeneration (for example, *rds*^-/-^ (Spencer et al., 2019), *Tmem138^-/-^* (Guo et al., 2022), *Spata7^-/-^* (Dharmat et al., 2018)), and *Rpgr^-/-^* (Megaw et al., 2024), however the degree of EV formation in comparison to disc formation differs among the different mouse models. Proteomic analyses of EVs in retinas deficient in CEP290 or in other proteins important for proper OS formation will be key in understanding the mechanisms of EV formation and providing insights into their possible roles in cargo-filtering and membrane protein sorting. Given that CEP290 loss does not affect NPHP5, a key TZ structural protein, or BBS4, a member of the BBSome trafficking complex, localization to the cilium, in contrast to findings demonstrating mislocalization of BBS4 and NPHP5 with loss of CEP290 (BBS4: (Kobayashi et al., 2014, Stowe et al., 2012, Klinger et al., 2014, Barbelanne et al., 2015); NPHP5: (Wiegering et al., 2021, Kim et al., 2018)), many questions remain about the mechanisms behind OS disc formation and protein trafficking in the photoreceptor cilium.

Although ciliopathy mutations in either *SPATA7* or *BBS4* cause blinding diseases through disruption of CC structure and function (Mykytyn et al., 2004, Wang et al., 2009, Katsanis et al., 2002), as CEP290 mutations do, the rates of degeneration and structural aberrations differ among them. Our results suggest that disruption of CC protein trafficking or localization by multiple mechanisms, including disruption of the distal CC through interruption of the luminal scaffold, may cause MT splaying. Because the ciliopathy mouse mutants studied here display varying degrees of OS disc disorganization, EVs present in the OS region, and axonemal MT splaying, it is possible that the membranous structures described in Fig. 7, within the lumen of their axonemal MTs at the distal CC/mid axoneme region, may be related to improperly forming discs and interruptions in IFT trafficking.

### Differential Roles of N- and C-terminal CEP290

CEP290^NN^ mutants retained sufficient N-terminal CEP290 at the PCC for normal CC formation (*i.e.*, with normal length and presence of ciliary membrane, Y-links, and ciliary necklace), in contrast to the CEP290^KO^. However, defective protein trafficking and outer segment (OS) disc formation, accompanied by extracellular vesicle (EV) buildup, were observed in both. Given that there is a difference in fluorescence patterns – punctate like staining in CEP290^KO^ but not in CEP290^NN^ – of certain OS proteins such as PRPH2 and PDE6β, it is possible that CC trafficking is also reliant on the proteins that are mislocalized in the CEP290^KO^ (AHI1 and NPHP1) but normally localized in the CEP290^NN^.

Interestingly, the truncation of the last two exons, located within the myosin-tail homology domain in CEP290 (CEP290^NN^), in mice results in phenotypes reminiscent of Joubert Syndrome (Datta et al., 2019). In contrast, the retinal degeneration and anosmic (McEwen et al., 2007) *Rd16* mouse model possesses an in-frame deletion of multiple exons within the C-terminal myosin-tail homology domain of CEP290, yet these mice do not display Joubert-like syndromic ciliary phenotypes. These results underscore the importance of the myosin-tail homology domain specifically in photoreceptor OS maintenance, and of the C-terminal end of this domain in ciliogenesis and MT stability within cilia throughout the body.

Although many canonical TZ proteins and structures localize throughout the length of the CC (Robichaux et al., 2019, Potter et al., 2021a, Rohlich, 1975), we have observed a distinct localization pattern for a subset of TZ proteins in the murine CC that localize proximally and not throughout the length of the CC. These include CEP78, CEP89, and NPHP8/RPGRIP1L (Nikopoulos et al., 2016). We have also documented a difference in spacing of ciliary necklace beads between the PCC region and the rest of the CC (Zhang et al., 2023). This proximal CC was largely unaffected in the CEP290^KO^ model, with Y-link and ciliary necklace structures preserved only in this region of the CC. PCC localization was retained for some CC proteins (AHI1 and NPHP1), as observed in the *Spata^-/-^* mice (Dharmat et al., 2018). Since TZ assembly occurs prior to axoneme extension (Insinna et al., 2019b, Insinna et al., 2019a, Lu et al., 2015), and since the sub-TZ appears largely normal in the absence of CEP290, we hypothesize that the assembly of the “sub-TZ” proteins and extension of the microtubules in photoreceptor cilia does not require full-length CEP290 function, although CEP290 appears to be important for the efficiency of this process.

This study highlights CEP290’s multifaceted role in photoreceptor ciliogenesis, CC stability, and protein trafficking, confirming that the critical roles of CEP290 in CC and OS formation emerge at early developmental stages, in addition to being essential in maintaining those structures at later stages, which will deepen our understanding of CC functions and their implications for ciliopathies.

## Methods

### Animals

All WT laboratory *Mus musculus* were C57BL/6 J between the ages of 10 days and 6 months.

Mice were kept on a 12-hour light/dark cycle. CEP290^NN^ (Cep290^tm1.1Jgg^/J; stock 013702) mice were obtained from The Jackson Laboratory. CEP290^KO^ (Rachel et al., 2015) mice were obtained from Anand Swaroop at NIH (KO). These animals are both global mutants, with homozygotes demonstrating many ciliary phenotypes such as male infertility, hydrocephalus, and higher rates of embryonic death. The *Bbs4^-/-^* mice were obtained from Dr. Samuel Wu and were originally characterized in (Eichers et al., 2006). *Spata7^-/-^* mice were a gift from Rui Chen (Baylor College of Medicine), and the mouse generation is detailed in (Eblimit et al., 2015). Retinas/eyecups from the CEP290 mutants and WT littermate controls, as well as the SPATA7^-/-^ mice were collected at post-natal day 10, and from *Bbs4^-/-^* collected at P30. Retina samples from at least 3 different mice per genotype were used for all immunoblot experiments. Eyecups from at least 3 different mice per genotype were used for sections for SIM, and each antibody condition was stained for on each *n* and imaged in at least 3 different areas. All STORM, expansion, and TEM conditions were repeated from multiple sections from at least 2 mice per genotype. Mice from both sexes were used indiscriminately, and as many mutant animals were used as possible, given that breeding large amounts of CEP290 mutant mice is difficult. All experimental procedures involving mice were approved by the Institutional Animal Care and Use Committee of Baylor College of Medicine.

### Human Subjects

This study adhered to the tenets of the Declaration of Helsinki and was approved by the Ethics Committees of Cantonal Committee of Canton Vaud for Research Activities on Human Subjects, the Ethikkommission Nordwest-und Zentralschweiz. Written informed consent was obtained from all individuals or their legal guardians prior to their inclusion in this study. Retinal tissue was collected following enucleation, and data was generated from the tissue of one subject.

#### Antibodies

The following primary antibodies were used: anti-centrin (Millipore Cat# 04-1624; 10ug for STORM/Expansion; 1:100 for IF), anti-INPP5e (ProteinTech Cat# 17797-1-AP; 10ug for STORM), anti-CEP290 (Bethyl Cat# A301-659A ; 2ug/ml for WB; 10ug for STORM/Expansion/ImmunoEM; 1:100 for IF), anti-CEP290 (ProteinTech Cat# 22490-1-AP; 10ug for STORM/Expansion/ImmunoEM), anti-CEP290 (BiCell Cat# 90006; 1ug/ml for WB; 5ug for STORM/Expansion/ImmunoEM; 1:50 for IF), anti-RP1 (gift from Eric Pierce, Harvard University; 1:1000 for IF), anti-RPGR (gift from Hemant Khanna; 10ug for STORM, 1:100 for IF), anti-acetylated tubulin (SantaCruz Cat# sc23950af647; 10ug for IF/STORM/Expansion), anti-GT335 (Adipogen Cat# AG-20B-0020-C100 (GT335); 1:100 for IF; 5ug for STORM/Expansion; 1:2000 for WB), anti-BBS4 (BiCell Cat# 90204; 1:100 for IF), anti-NPHP1 (BiCell Cat# 90001; 1:100 for IF), anti-CEP89 (BiCell Cat# 01079; 1:100 for IF), anti-CP110 (ProteinTech Cat# 12780-1-AP; 1:100 for IF), anti-CEP97 (ProteinTech Cat# 22050-1-AP; 1:100 for IF), anti-AHI1 (ProteinTech Cat# 22045; 1:100 for IF), anti-NPHP8 (BiCell Cat# 90008; 1:100 for IF), anti-IFT88 (ProteinTech Cat# 13967-1-AP; 1:100 for IF), anti-IFT140 (Gift from Gregory Pazour, UMass; 1:100 for IF), anti-IFT81 (ProteinTech Cat# 11744, 1:100 for IF), anti-Rootletin (SantaCruz Cat# sc-67824; 1:500 for IF), anti-GAPDH (Fitzgerald Cat# 10R-G109a; 1:20,000 for WB), anti-β actin (CST Cat# 3700S; 1:1,000 for WB), anti-PDE6β (SantaCruz Cat# sc-377486; 1:500 for WB; 1:250 for IF), anti-pde6α (ABR Cat# PA1-720; 2ug for WB), anti-PRPH2 (gift from Andrew Goldberg, Oakland University; 1:3000 for WB; 1:500 for IF), anti-CNGA1 (EMD Cat# MABN2617-100UG; 1:100 for IF), anti-arl13b (Proteintech Cat# 17711-1-AP; 1.4ug for WB; 1:250 for IF), anti-cArr3 (Sigma Cat# AB15282; 1:1000 for IF), anti-STX3 (Proteintech Cat# 15556-1-AP; 1ug for WB; 1:500 for IF), anti-Rho-C-1D4 (Millipore Cat# MAB5356; 1:1000 for IF), anti-ROM1 (Proteintech Cat# 21984-1-AP; 1:500 for WB; 1:250 for IF), anti-GC1 (Santa Cruz Cat# sc-376217; 1:150 for WB; 1:500 for IF). The following primary antibodies were used: PNA (Vector Laboratories Cat# RL-1072; 1:1000 for IF), F(ab’)2-goat anti-mouse IgG Alexa 647 (Thermo Fisher Scientific Cat# A48289TR), F(ab’)2-goat anti-mouse IgG Alexa 555 (Thermo Fisher Scientific Cat# A21425), F(ab’)2-goat anti-rabbit IgG Alexa 647 (Thermo Fisher Scientific Cat# A-21246), F(ab’)2-goat anti-rabbit IgG Alexa 555 (Thermo Fisher Scientific Cat# A-21430), F(ab’)2-goat anti-rabbit IgG Alexa 488 (Thermo Fisher Scientific Cat# A-11070), F(ab’)2-goat anti-mouse IgG CF568 (Biotium Cat# 20109), F(ab’)2-goat anti-rabbit IgG CF568 (Biotium Cat# 20099), all used at 1:1000 for IF, 7ug for STORM/Expansion. Nanogold-Fab’ goat anti-rabbit (Nanoprobes Cat# 2004), 15ul were used for ImmunoGoldEM.

#### Immunofluorescence

For confocal/Airyscan/SIM immunofluorescence, mouse eyes were enucleated, cornea and lens dissected out, and either immediately frozen in OCT (Moye et al., 2023, Potter et al., 2021b) for ciliary staining, or fixed in 4% PFA at RT for 2 hours. The fixed eyecups were then incubated in 30% sucrose overnight at 4C, then in a 1:1 mix of 30% sucrose and optical cutting temperature medium (OCT) before being frozen in OCT. 8µm sections were collected on Superfrost+ slides (VWR, Cat# 48311-703, Radnor, Pennsylvania, USA). Unfixed sections (for ciliary staining) were fixed with 4% PFA (in 1xPBS) for 2 min prior to immunolabeling. For immunolabeling, sections were quenched with 100 mM glycine (in 1x PBS) for 10 min at RT. Sections were then incubated with blocking solution: 10% normal goat serum (NGS) (VWR #102038-714, Radnor, Pennsylvania, USA), 0.2% Triton X-100, 2% Fish Skin Gelatin (Sigma, Cat# G7041, Burlington, Massachusetts, USA), in 1x PBS, for 1 h at RT. Primary antibodies were made in the same block buffer (1µg – 5µg) and sections were incubated overnight at 4°C in a humidified chamber. The next day, after washing 3 times in 1xPBS, 5 min each, sections were incubated with 1 µg of fluorescent secondary antibodies (again in block buffer) for 1 h at RT. After washing, sections were mounted with ProLong Glass Antifade Mountant (Thermo Fisher Scientific Cat# P36980, Waltham, Massachusetts, USA).

For STORM and expansion immunofluorescence, dissected mouse retinas were dissected in ice cold buffered Ames’ media (Sigma, Cat# A1420, Burlington, Massachusetts, USA) and were fixed in 4% PFA diluted in Ames’ for 5 min on ice (for whole retina samples). Retinas were quenched in 100 mM glycine for 30 min at RT, then incubated in 1 mL of SUPER block solution: 15% NGS, 5% bovine serum albumin (BSA) (Sigma, Cat# B6917, Burlington, Massachusetts, USA) + 0.5% BSA-c (Aurion, VWR, Cat# 25557, Radnor, Pennsylvania, USA) + 2% fish skin gelatin (Sigma, Cat# G7041, Burlington, Massachusetts, USA) + 0.05% saponin (Thermo Fisher, Cat# A1882022, Waltham, Massachusetts, USA) + 1x protease inhibitor cocktail (GenDepot, Cat# P3100-005, Katy, Texas, USA), in low-adhesion microcentrifuge tubes (VWR, Cat# 49003-230, Radnor, Pennsylvania, USA) for 3 h at 4°C. 1 µg – 5 µg of primary antibodies were added directly to the blocking solution at 4°C and left to incubate for 3 days with mild agitation. Retinas were washed 6 times for 10 min each in 2% NGS diluted in Ames’ prior to probing with 4 µg – 8 µg of secondary antibodies diluted in 1 ml of 2% NGS in Ames’ + 1x protease inhibitor cocktail at 4°C overnight with mild agitation. Retinas were then washed 6 times, 5 minutes each in 2% NGS diluted in Ames’.

For STORM, retinas were fixed in 2% PFA + 0.5% glutaraldehyde diluted in 1xPBS for 30 min at 4°C with mild agitation. They were then dehydrated in an ethanol series (50%, 70%, 90%, 100%, 100%) for 15 min each in half dram vials on a RT roller. Dehydrated retinas were then embedded in Ultra Bed Low Viscosity Epoxy resin (EMS Cat# 14310, Hatfield, Pennsylvania, USA) as outlined previously (Robichaux et al., 2019). A Leica UCT ultramicrotome and glass knives were used to make 0.5 µm – 1 µm thin retinal cross sections that were placed onto 35mm glass-bottom dishes with a 10mm microwell (MatTek Life Sciences, Cat# P35G-1.5-14-C, Ashland, Massachusetts, USA), and chemically etched in a mild sodium ethoxide solution (∼1% diluted in pure ethanol for 0.5 – 1.5 714 h) as previously described (Robichaux et al., 2019). Immediately prior to STORM imaging, etched sections were mounted in a STORM imaging buffer adapted from (Albrecht et al., 2022): 50 mM Tris (pH 716 8.0), 10 mM NaCl, 10 mM sodium sulfite, 10% glucose, 40 mM cysteamine hydrochloride (MEA, Chem Impex/VWR, Cat# 102574-806, Radnor, Pennsylvania, USA), 143 mM BME, and 1 mM cyclooctatetraene (Sigma Cat# 138924, Burlington, Massachusetts, USA), under a #1.5 glass coverslip that was sealed with quick-set epoxy resin (Devcon).

For expansion in mouse retina, the protocol is outlined in detail in (Moye et al., 2023), with an expansion factor of 4. Briefly, mouse retinas were stained according to the STORM protocol above. After crosslinking to Acryloyl-X SE (Life Tech, A20770, Carlsbad, California, USA), they were gelled in a polyacrylamide solution and subjected to denaturation with high salt-high heat. The gels were then re-stained in the primary and secondary antibodies prior to freezing in O.C.T. compound. 10-20 μm sections were cut on a cryostat and then expanded in a beaker of di-water. These expanded sections were then placed on glass slides and covered with a #1.5 glass coverslip, mounted in water, for immediate imaging.

For iterative expansion microscopy in human retina, the gelation, staining, and expansion was performed just as laid out in (Louvel et al., 2023), the only changes pertained to antibodies/dilutions, listed in Table 1. An expansion factor of ∼14x was obtained (Supplemental Figure S2B), by taking the FWHM of the BB in rods (average of 3.2 ± 190 µm) and dividing by 230 nm (assuming that the width of BB in photoreceptors is consistent between mouse and human).

#### Imaging

**Confocal** scanning was performed on a Zeiss LSM 710 using a 63x/NA 1.4 oil objective. Airyscan images were acquired using a Zeiss LSM 880 using a 63x/NA 1.4 oil objective. **SIM** imaging was performed on a DeltaVision OMX Blaze v4 (GE Healthcare, now Cytiva) equipped with 405 nm, 488 nm, 568 nm, and 647 nm lasers and a BGR filter drawer; a PLANPON6 60×/NA 1.42 (Olympus) using oil with a refractive index of 1.520; and front illuminated Edge sCMOS (PCO). For imaging expanded tissue on the OMX SIM, the only change was use of a 2.52 refractive index immersion oil to try to match water/thick tissue as much as possible. All **STORM** acquisitions were performed at RT on a Nikon N-STORM 5.0 system equipped with an Andor iXON Ultra DU-897U ENCCD camera with a SR HP Apochromat TIRF (total internal reflection fluorescence) 100x/NA 1.49 oil immersion objective. The full system details and STORM acquisition protocol are outlined in (Potter et al., 2021b, Robichaux et al., 2022). **iUExM** Imaging was performed on 35mm glass bottom dishes with a 10mm microwell (MatTek Life Sciences, Cat# P35G-1.5-14-C, Ashland, Massachusetts, USA) that had been coated in Poly-L-lysine. A gel slice was placed on the dish, a drop of water added, and coverslip added on top. The imaging was performed on a Leica Stellaris 8 Falcon using HyD lasers and a 40x HC PL APO CORR CS2 water immersion objective, NA 1.10, often with an optical zoom between 2-5.

#### TEM

Retinas for TEM were immediately fixed in 2% PFA (Fisher Scientific # 50980487, Waltham, Massachusetts, USA) + 2% glutaraldehyde (Fisher Scientific # 5026218, Waltham, Massachusetts, USA + 4.5 mM CaCl2 in 50 mM MOPS buffer (pH 7.4) for 2-5 h at 4°C on a roller. Retinas were then subjected to the exact same protocol as performed in (Potter et al., 2021b) (vibratome slices) and (Robichaux et al., 2022) (full retina). 70 nm ultramicrotome sections were cut from the resin blocks using a Diatome Ultra 45° diamond knife and collected onto copper slot-grids (VWR, Cat# 102100-816, Radnor, Pennsylvania, USA). Grids were post-stained in 1.2% uranyl acetate diluted in water for 4 min, rinsed 6 times in water, and allowed to completely dry before staining with a lead citrate solution (EMS, #22410, Hatfield, Pennsylvania, USA) for 4 min. Grids were then rinsed in water and dried overnight. Grids were imaged on either a JEOL 1400 Plus electron microscope with an AMT XR-16 mid-mount 16-megapixel digital camera or on a JEOL JEM-1400Flash 120 kV TEM with a high-contrast pole piece and a 15 megapixel AMT NanoSprint15 sCMOS camera. For each microscope, AMT software was used for image acquisition and images were subsequently cropped with slight contrast adjustments in FIJI//ImageJ (Schneider et al., 2012).

#### Immuno-EM

Retinas were dissected immunolabeled as described in the STORM section except the pre-fixation solution was 4% PFA + 2.5% glutaraldehyde in Ames’, nanogold-conjugated secondaries were used (5 - 7.5 µg), and the post-fixation solution was 2% PFA + 2% glutaraldehyde + 4.5 mM CaCl2 in 50 mM MOPS buffer (pH 7.4). Retinas were then rinsed with water and enhanced using HQ Silver Kit (Nanoprobes, Cat# 2012) reagents in half dram vials for 4 min at RT with agitation. Enhanced retinas were then immediately rinsed with water, incubated in 1% tannic acid + 0.5% glutaraldehyde in 0.1 M HEPES (pH 7.5) for 1 h on a RT roller, rinsed with water, incubated 1% uranyl acetate in 0.1 M maleate buffer (pH 6.0) for 1 h on a room temperature roller, and rinsed a final time with water. Retinas were ethanol dehydrated, resin embedded in Eponate resin, sections were cut, and grids were stained and imaged as outlined in the TEM section. Grids imaged as outlined above.

#### Image processing

SIM images were were reconstructed using SI reconstruction and OMX alignment in Softworx 7 software using default settings. STORM reconstruction data were processed in NIS Elements Ar v5.30.05 (Nikon) using the N-STORM Analysis modules. After analysis, SIM and STORM reconstructions were processed in Fiji/ImageJ (Schindelin et al., 2012). The Straighten tool was applied to straighten individual curved or bent cilia for both SIM and STORM to acquire accurate profiles. Airyscan images first went through “Airyscan processing” in ZenBlue software before being exported into Fiji/ImageJ. iUExM images underwent Lightning processing on the Leica Stellaris 8 Falcon immediately following image capture. FIJI was used for image visualization and basic adjustments of all SIM, STORM, TEM, and confocal imaging.

#### Localization analysis

Performed in Fiji/ImageJ. TEM images of longitudinal CC were first thresholded for better visualization of the SEGCs. For radial angle measurements, a circle was first drawn around near-circular CC (oblong CC discarded) and that circle was copied onto the thresholded version of the image. The x,y coordinates of the center of the circle and of the SEGCs (Analyze Particles size 2-Infinity, circularity 0-1) were then used to calculate the radial distances of each SEGC (or DMT) from the centroid of the CC. These radii distributions were plotted as histograms in Prism.

#### Western Blotting

For retinal lysate western blotting, mouse retinas were dissected into RIPA buffer with Protease Inhibitor cocktail (Roche, Cat# 11836153001, Basel, Switzerland). Lysis was performed through sonication and samples were loaded onto either a 10% Tris-Glycine or a 3%-8% Tris-Acetate gel (for CEP290). Either a Precision Plus Dual Color ladder (Bio-Rad, Cat# 1610374, Hercules, California, USA) or the HiMark Protein ladder (BioRad Cat# LC5699, Hercules, California, USA) were used. Corresponding buffers (Tris-Glycine or Tris-Acetate) for SDS-PAGE and membrane transfer (onto ImmobilonFL Transfer Membrane PVDF (pore size: 0.45 µm) (LI-COR Cat# 92760001, Lincoln, Nebraska, USA) were used depending on which gel was being run. Membranes were subsequently blocked using Intercept Blocking Buffer (LI-COR, Cat# 927-6000, Lincoln, Nebraska, USA) for 1 h, then incubated with primary antibody in Blocking Buffer + 0.2% Tween-20 (antibodies were diluted at 1:500 - 1:5000) overnight at 4°C, except for GAPDH (1 hour incubation at 1:20,000). Membranes were washed in 1xPBS + 0.1% Tween-20 (PBS-T) 3 times, 5 min each before secondary staining with LiCor 800CW or 680RD secondary antibodies (1:5,000 each) diluted in Blocking Buffer + 0.2% Tween-20 for 1 h. Membranes were washed then imaged for fluorescence on an Azure scanner using both 800 and 680 channels. Images were analyzed on Azure software.

#### Statistical analysis

Frequencies were calculated for each ciliogenesis stage in Figure 4, and T-tests were performed to compare between genotypes. Fishers Exact Test (Supplemental Figure S4) were used for population comparisons. T-tests were also used for quantitative analysis of centrin lengths.

## ACKNOWLEDGEMENTS

Thanks to the BCM OiVM and Imaging Core Facilities where Airyscan and SIM imaging experiments were performed by ARM. Thanks to the Robichaux lab at West Virginia University for performing some of the STORM imaging. The authors also thank the TEM facility at Rice University and Lita Duraine at the Neuroscience Research Institute for TEM imaging assistance and expert advice on sample preparation and sectioning. We thank the Imaging Core facility (IMCF, Biozentrum, University of Basel) and in particular Alexia Loynton-Ferrand for the technical assistance provided on the Stellaris 8 Falcon microscope. The authors declare no competing financial interests.

## AUTHOR CONTRIBUTIONS

Conceptualization: ARM, TW; Investigation: ARM, MAR; Resources: TW, MAR, CR, APM; Writing – original draft: ARM; Writing – reviewing & editing: MAR, MAA, TW; Data analysis input: MAR, MAA, TW; Visualization: ARM.

## Acronyms

MT: microtubules
DMT: doublet microtubules
CC: connecting cilium
PCC/DCC: proximal/distal connecting cilium
OS: outer segment
IS: inner segment
BB: basal body
TZ: transition zone
IFT: intraflagellar transport
TEM: transmission electron microscopy
STORM: stochastic optical reconstruction microscopy
SIM: structured illumination microscopy

