## Supplemental Figures for "Ciliopathy-associated protein, CEP290, is required for ciliary necklace and outer segment membrane formation in retinal photoreceptors"

**Supplemental Figures and Legends**

**Movie 1. CEP290 localization in human cone photoreceptors.** Iterative expansion microscopy (iUExM) confocal images from expanded human retina of photoreceptor cilia immunostained for tubulin (cyan) and CEP290 N-terminus. DAPI (grey) is used to display non-specific membrane and rootlet staining.

**Movie 2. CEP290 localization in human rod photoreceptors.** Iterative expansion microscopy (iUExM) confocal images from expanded human retina of photoreceptor cilia immunostained for tubulin (cyan) and CEP290 N-terminus, from a cross-sectional view. DAPI (grey) is used to display non-specific membrane and rootlet staining.

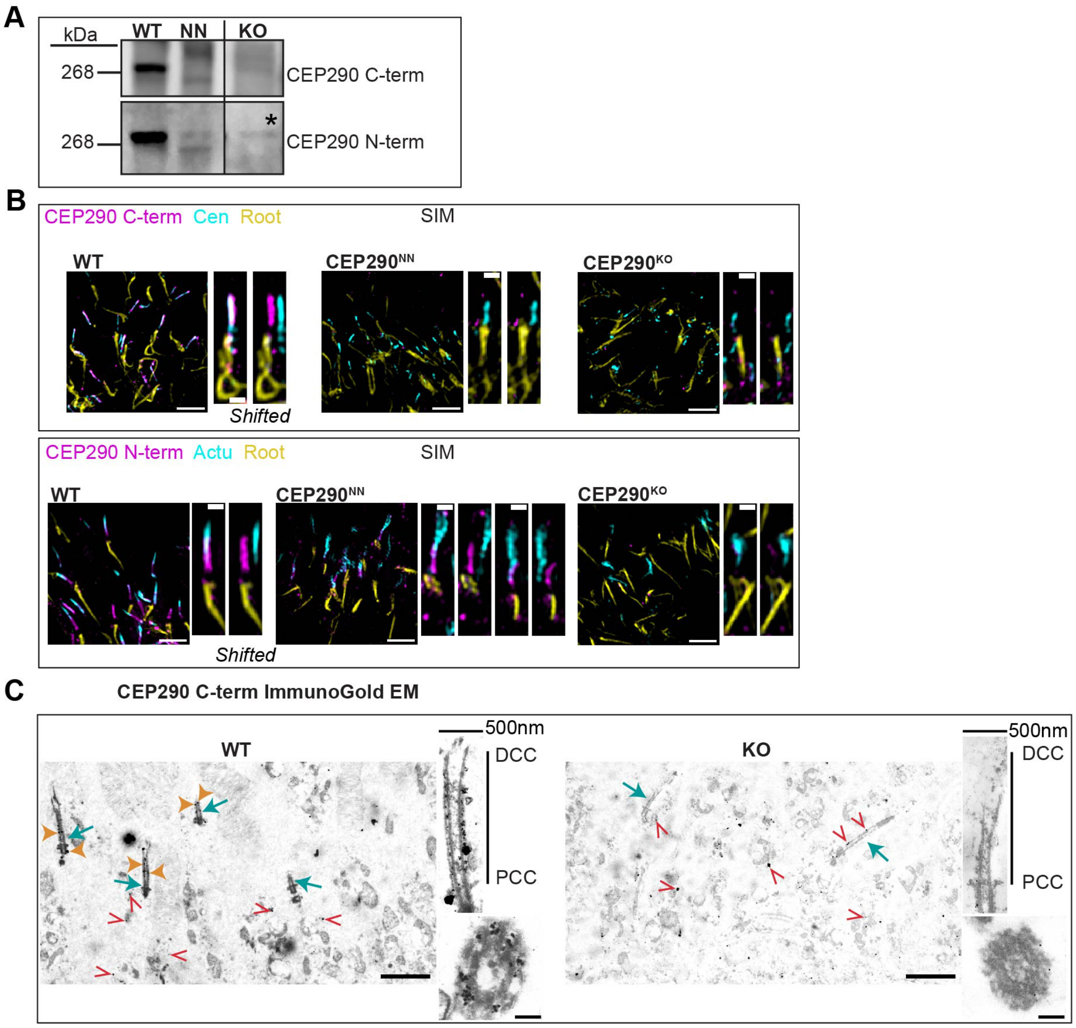


**Supplemental Figure S1. CEP290 Antibody Validation.** (A) Western blot analysis of P10 retinal lysates from WT, $CEP290^{NN}$, and $CEP290^{KO}$ probed with the C-terminal antibody (Bethyl labs) and an N-terminal antibody (BiCell). Clear bands at ~290kDa were observed in the WT, with no bands in the $CEP290^{KO}$ (there was a faint band above the WT band, which is also observed in lanes with no sample loaded, indicating it as a non-specific band). In $CEP290^{NN}$, there are some lower molecular weight bands observed, consistent with previous reports [64]. (B) Immunofluorescence (imaged with SIM) was performed on P10 retinas from WT and CEP290 mutants, with no CEP290 staining in the KO, or the $CEP290^{NN}$ with C-term antibody (SIM). Notably, the N-terminal antibody detects CEP290 at the base of the CC in the $CEP290^{NN}$. (C) Immunogold EM at P10 in WT and $CEP290^{KO}$ (blue arrows indicate cilia, orange arrowheads indicate CEP290 staining, red open arrowheads indicate non-specific nanogolds). In the $CEP290^{KO}$, there is no specific staining. ImmunoGold EM = 2 $\mu$m (left), 500 nm (width of longitudinal CC), and 100nm (transverse section); SIM = 2 $\mu$m, 500 nm zoomed images. Each length from proximal CC (PCC) to distal CC (DCC) is 1.5 $\mu$m.

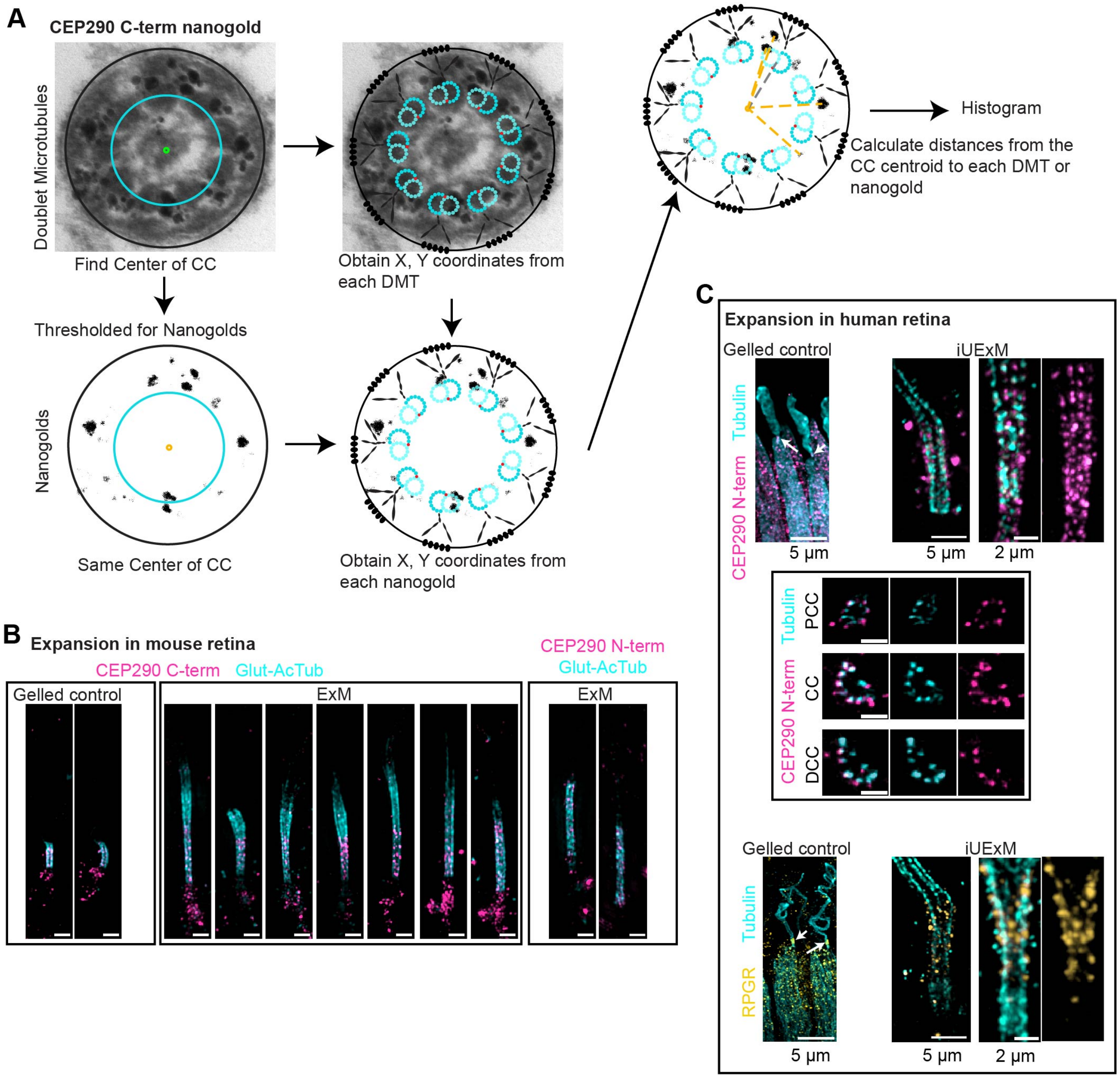


**Supplemental Figure S2. Localization analysis for CEP290.** (A) ImmunoEM transverse images of WT retina stained with C-terminal CEP290 antibody with a diagram to demonstrate how radial distributions were obtained. Transverse ImmunoEM images were thresholded, and the X, Y coordinates were plotted in comparison to the center of each cilium and the radii from the centroid to each SEGC was obtained. A histogram was then created showing the distribution in radial distances. (B) SIM z-projection images of single expanded CC stained for either CEP290 C-terminus or CEP290 N-terminus with a glutamylated/acetylated tubulin mix (cyan) as the microtubule marker from expansion experiments in mice. Scale bar = 1000 nm. (C) Gelled control images from the iUExM experiments, arrows point to the CC regions. To the right are further examples of iUExM images not shown in Figure 1.

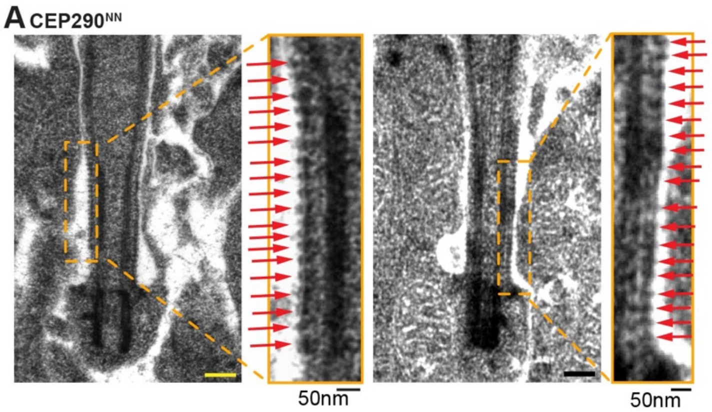


**Supplemental Figure S3. CEP290$^{NN}$ CC membrane similar to WT.** (A) Longitudinal TEM images of P10 eyecups from CEP290$^{NN}$ retina. Left, image depicting CC with ciliary necklace (membrane beads); right, zoomed images with ciliary membrane protrusions marked with red arrows.

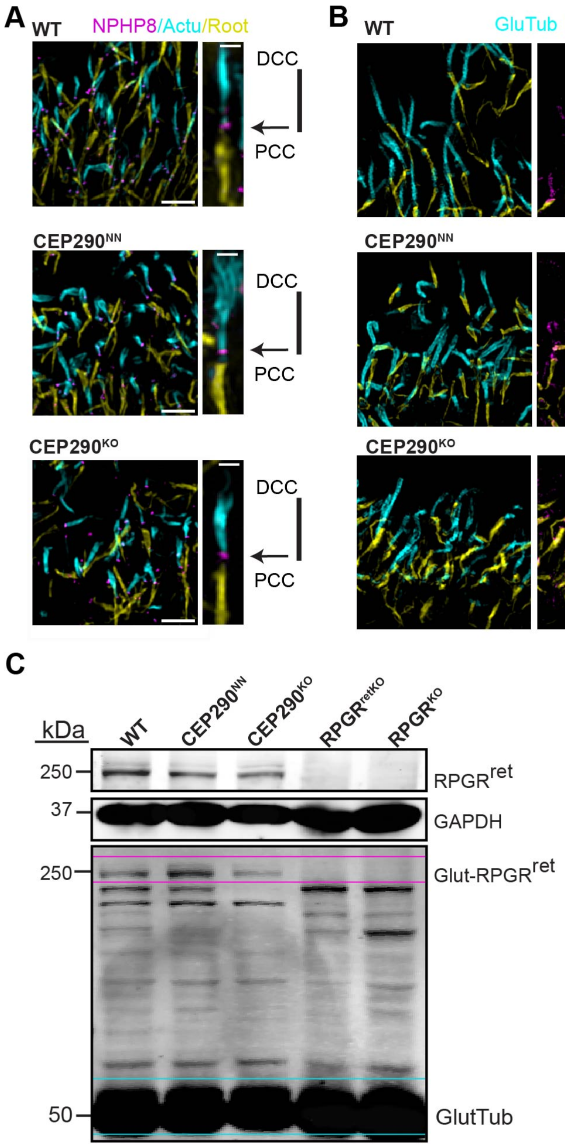


**Supplemental Figure S4. Variation of Transition Zone protein localization in photoreceptor CC.** (A) SIM low magnification images (left) of cilia from WT and CEP290 mutant retinal cryosections at P10, displaying the localization of NPHP8/RPGRIP1L, with proximal CC (PCC) and distal CC (DCC) marked. Scale bar = 2 $\mu$m. SIM (right) images of a representative cilium. Scale bar = 500 nm. (B) SIM images demonstrating the variability of RPGR mislocalization in CEP290 mutant photoreceptors, from absence of ciliary localization, to random distribution throughout the CC. (C) Immunoblot analysis of retinal lysates from WT, CEP290$^{NN}$, and CEP290$^{KO}$ (P10) as well as an RPGR KO and an RPGR retina-specific isoform

KO (adult) probed for RPGR and polyglutamyl, which is also a marker for the retina-specific isoform of RPGR showing RPGR protein is expressed in the CEP290 mutant retina.

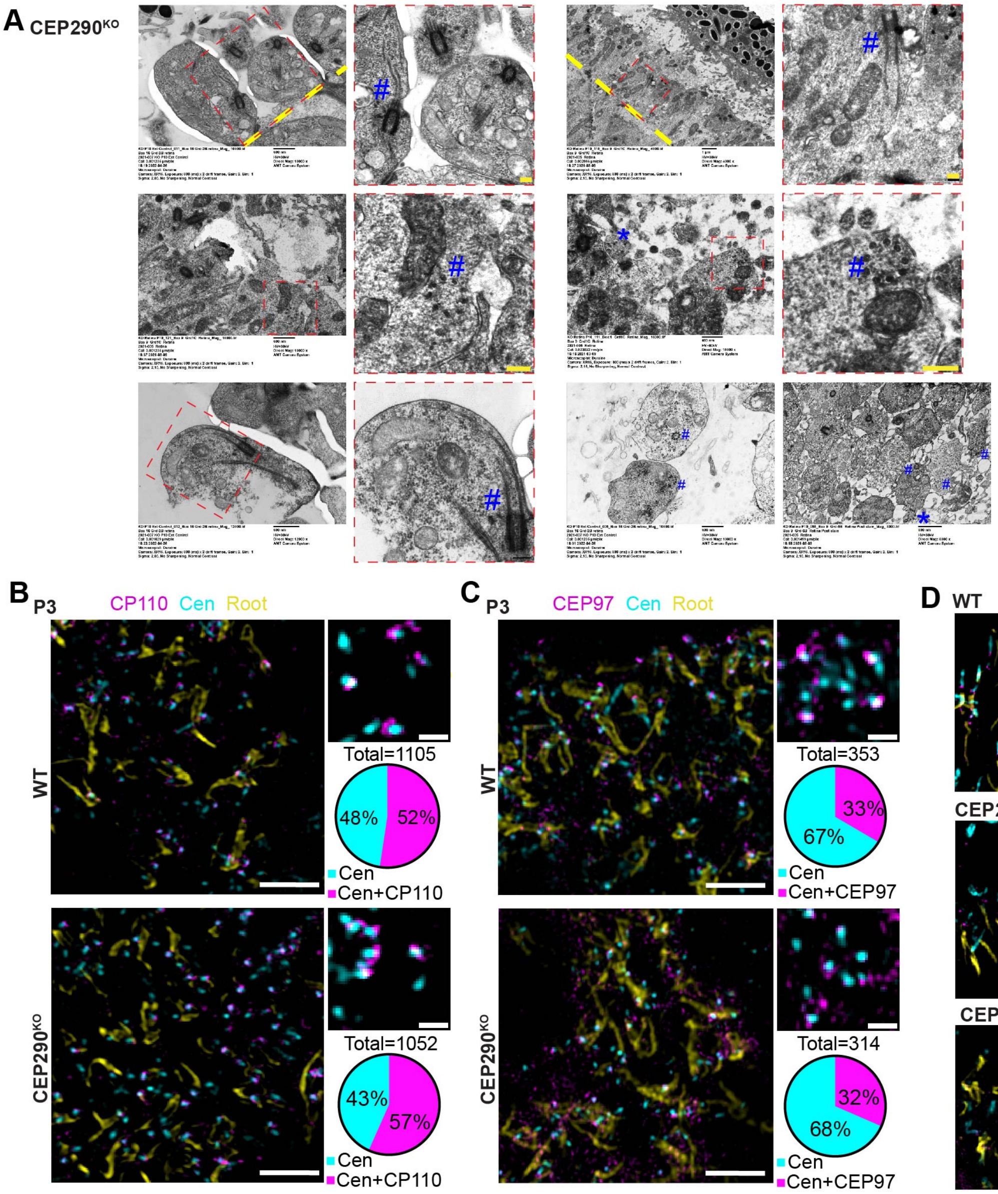


**Supplemental Figure S5. Ciliogenesis at P3 similar between WT and CEP290$^{KO}$.** (A) Zoomed out electron micrographs used in Figure 3, as well as two additional images displaying microtubules inside cytoplasm in the CEP290$^{KO}$ retina at P10. The areas used for figure 3 are boxed in red dashed lines. (B, C) SIM images of P3 retinal cryosections from WT and CEP290$^{KO}$ mice stained with indicated markers of ciliogenesis. Scale bars = 2 $\mu$m, zoomed insets = 500 nm. Below each inset is a pie chart displaying the percentage of centrin labelled puncta either alone or associated with CP110 (B) or CEP97 (C) labelling. n=1 animal for each

WT and KO, measurements from multiple images taken from 3 individual sections. As such, there is no difference in the percentages between WT and $CEP290^{KO}$. (D) SIM low magnification images (left) of cilia from WT and CEP290 mutant retinal cryosections at P10, displaying the localization of CEP89, with proximal CC (PCC) and distal CC (DCC) marked. Scale bar = 2 $\mu$m. SIM (right) images of a representative cilium. Scale bar = 500 nm.

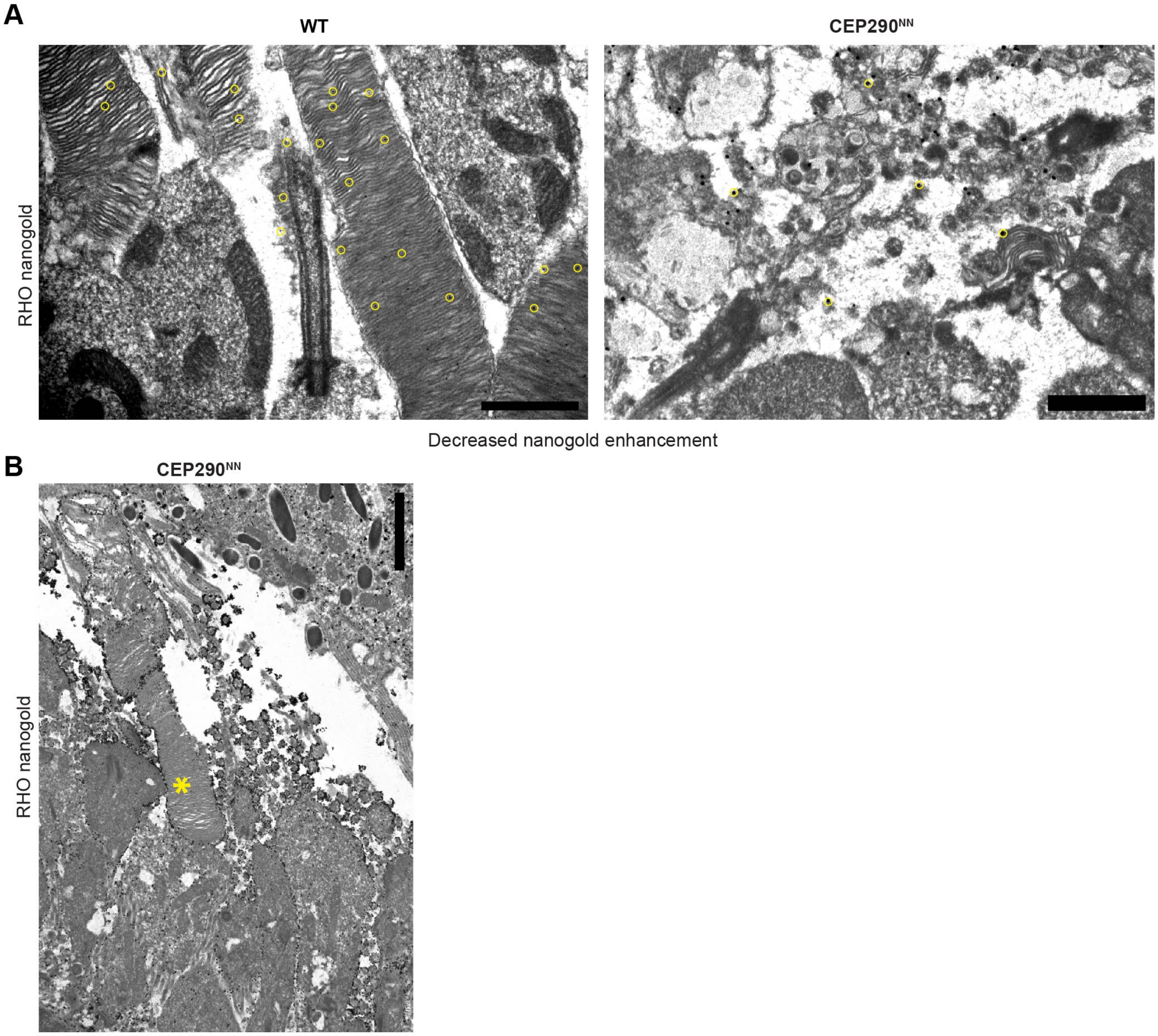


**Supplemental Figure S6. ImmunogoldEM control images from murine retina.** Immunogold labeling of Rhodopsin (1D4) in TEM images from WT and $CEP290^{NN}$ retinas at P10. (A) shows staining where nanogold enhancement was decreased to confirm specificity of the antibody labeling (yellow circles denote some examples of the golds). 1D4 labels OS discs in WT and vesicles in CEP290$^{NN}$. Scale bars = 1 $\mu$m. (B) Another example of immunogold labelling in the CEP290$^{NN}$ retinas that contain some OS discs. Scale bar = 2 $\mu$m.

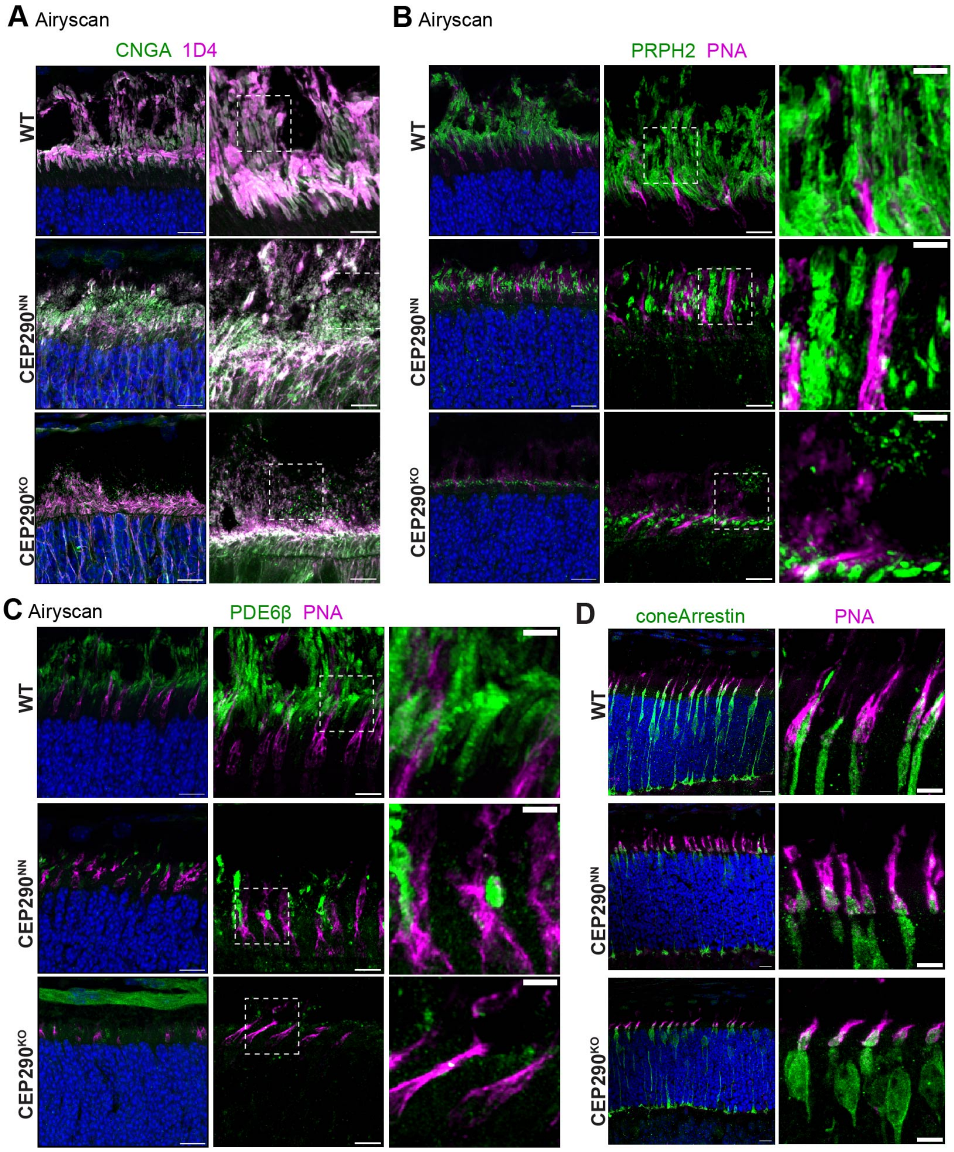


**Supplemental Figure S7. Outer segment protein localization in WT and CEP290 mutant retinas.** (A-D) Airyscan and confocal (D) images of fixed retinal cryosections at P10 from WT, CEP290$^{NN}$, and CEP290$^{KO}$ mice, stained with the indicated antibodies. Zoomed outsets are marked by dotted white boxes. (A) displays zoomed out examples from Figure 8D. PNA (peanut agglutinin) and cone Arrestin staining indicate cone numbers are comparable between WT and CEP290 mutants. Scale bar = 2 $\mu$m, with 200 nm scales marked to the side.

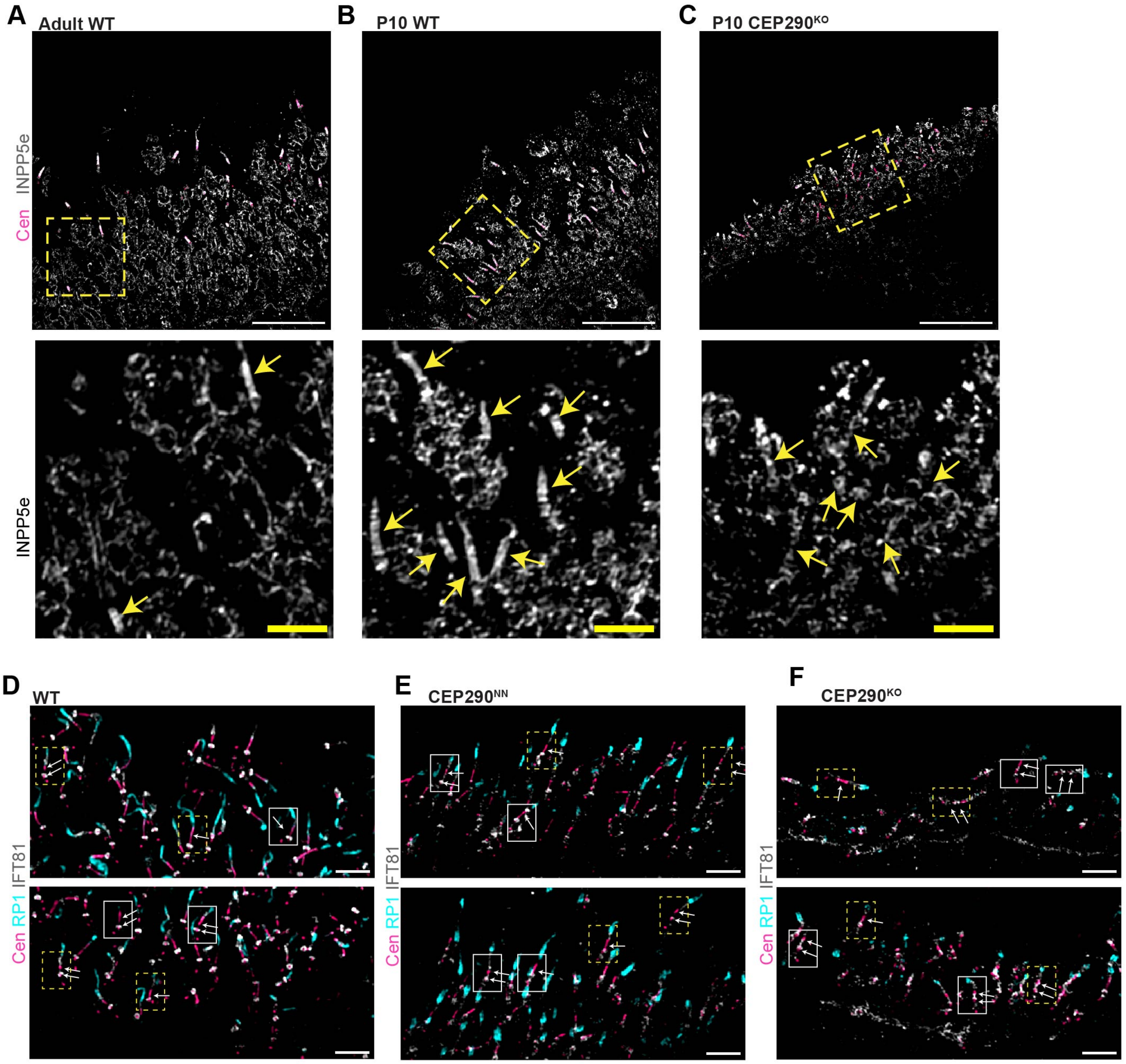


**Supplemental Figure S8. CC labeling in WT and CEP290 mutant retinas.** (A-C) SIM images showing INPP5E staining membranous structures within the IS (possibly mitochondrial membranes) as well as CC membranes and possibly into OS axonemes. Yellow dashed boxes show the regions which used in Figure 4 A, B. Below are also the zoomed insets, but with no centrin-labeling, CC indicated with arrows. Scale bar = 10 $\mu$m and 2 $\mu$m (below). (D-F) SIM images displaying localization of Centrin (pink), RP1 (cyan), and IFT81 (gray) in retinas of WT (A), CEP290$^{NN}$ (B), and CEP290$^{KO}$ (C) retinas at P10. Highlighted examples of full-length centrin and punctate centrin staining are boxed with arrows. Yellow dashed boxes are the zoomed insets used in Figure 4G. Scale bar = 2 $\mu$m.

**Blot Transparency.** Full Western Blots used in Supplemental Figures S1 and S4.

Supplementary Figure S1

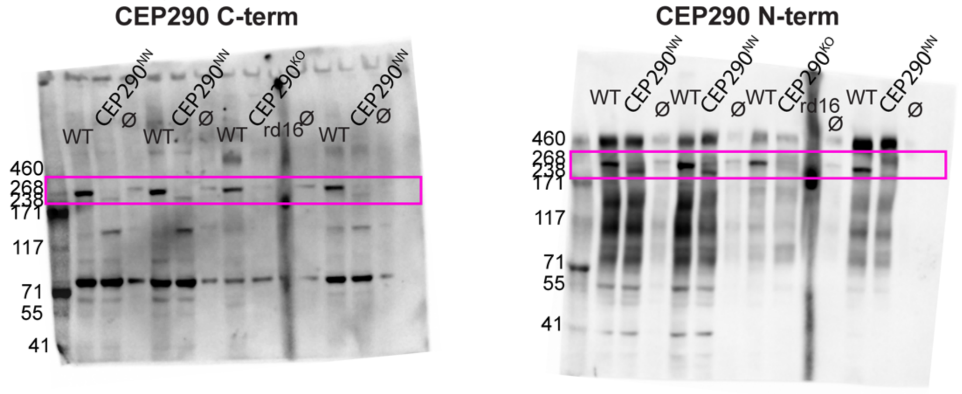


Supplementary Figure S4

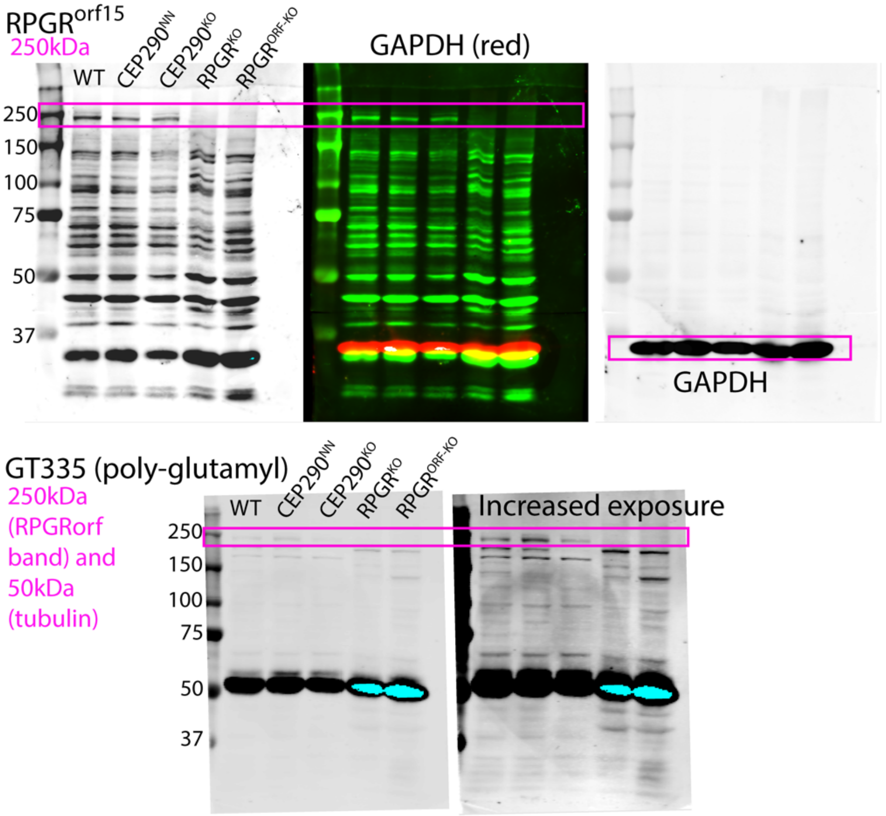
